# *Saccharomyces cerevisiae* H2A copies differentially contribute to recombination and CAG/CTG repeat maintenance, with a role for H2A.1 threonine 126

**DOI:** 10.1101/331413

**Authors:** Nealia C.M. House, Erica J. Polleys, Ishtiaque Quasem, Cailin E. Joyce, Oliver Takacsi-Nagy, Ryan B. Hayman, Jocelyn E. Krebs, Stephen M. Fuchs, Catherine H. Freudenreich

## Abstract

DNA are sites of genomic instability. Long CAG/CTG repeats form hairpin structures, are fragile, and can expand during DNA repair. The chromatin response to DNA damage can influence repair fidelity, but the knowledge of chromatin modifications involved in maintaining repair fidelity within repetitive DNA is limited. In a screen for CAG repeat fragility in *Saccharomyces cerevisiae*, histone 2A copy 1 (H2A.1) was identified to protect the repeat from increased rates of breakage. To address the role of H2A in CAG repeat instability, we tested the effect of deleting each histone H2 subytpe. Whereas deletion of *HTA2, HTZ1, HTB1*, and *HTB2* did not significantly affect CAG repeat maintenance, deletion of *HTA1* resulted in increased expansion frequency. Notably, mutation of threonine 126, unique to H2A.1, to a non-phosphorylatable alanine increased CAG repeat instability to a similar level as the *hta1Δ* mutant. CAG instability in the absence of HTA1 or mutation to *hta1-T126A* was dependent on the presence of the homologous recombination (HR) repair proteins Rad51, Rad52, and Rad57, and the Polδ subunit Pol32. In addition, sister chromatid recombination (SCR) was suppressed in the *hta1Δ* and *hta1-T126A* mutants and this suppression was epistatic to *pol32Δ.* Finally, break-induced replication (BIR) is impaired in the *hta1Δ* mutant, resulting in an altered repair profile. These data reveal differential roles for the H2A subtypes in DNA repair and implicate a new role for H2A.1 threonine-126 phosphorylation in mediating fidelity during HR repair and promoting SCR. Using a fragile, repetive DNA element to model endogenous DNA damage, our results demonstrate that H2A.1 plays a greater role than H2A.2 in promoting homology-dependent repair, suggesting H2A.1 is the true homolog of mammalian H2AX, whereas H2A.2 is functionally equivalent to mammalian H2A.

**Author Summary:** CAG/CTG trinuncleotide repeats are fragile sequences that when expanded can cause human disease. To evaluate the role of *S. cerevisiae* histone H2A copies in DNA repair, we have measured instability of an expanded CAG/CTG repeat tract and repair outcomes in H2A mutants. Although the two copies of H2A are nearly identical in amino acid sequence, we found that the CAG repeat is more unstable in the absence of H2A copy 1 (H2A.1) than H2A copy 2, and that this role appears to be partially dependent on a phosphorylatable threonine at residue 126 in the C-terminal tail of H2A.1. Further, we show through a series of genetic assays that H2A.1 plays a role in promoting homologous recombination events, including sister chromatid recombination and break-induced replication. Our results uncover a role for H2A.1 in mediating fidelity of repair within repetitive DNA, and demonstrate that modification of its unique Thr126 residue plays a role in regulating SCR. Given the dependence of HR repair on H2A.1 but not H2A.2, we conclude that H2A.1 plays a greater repair-specific role in the cell and therefore would be the true homolog of mammalian H2AX.

## Introduction

Integral to the eukaryotic DNA damage response is modification of the chromatin structure surrounding the break site (reviewed in (1-3)). In response to DNA double strand breaks (DSBs) and stalled or collapsed replication forks, the SQEL motif in the H2A C-terminal tail is phosphorylated at Ser129 (H2AX-Ser139 in mammals) by the Phosphoinositide 3-Kinase-Related Kinases (PIKKs), Mec1 and Tel1 (4, 5). This modification, termed γH2A (γH2AX in mammals), marks the site of damage and is propagated along the chromatin, detectable up to 50kb from the break site in yeast (6) and megabases in mammalian cells (7).

γH2A/γH2AX is the first detectable histone modification upon induction of a DSB, occurring within minutes after damage (6-8), and is required to initiate the cascade of histone modifications, chromatin remodeling, and repair factor recruitment and retention that is necessary for repair of the lesion. However, other modifiable residues in the H2A tails can also modulate repair factor binding and therefore contribute to repair: H2A/H2AX K5ac, K36ac, K119ub, and Y142ph are all documented to occur at sites of DSB repair (reviewed in (9)).

Whereas in humans over a dozen H2A variants have been identified (10, 11), just two copies of H2A exist in *S. cerevisiae*, encoded by the *HTA1* and *HTA2* genes (Figure 1A). The two copies of H2A are nearly identical in amino acid sequence except for a direct alanine-threonine switch in the C-terminal tail (Figure 1B); the underlying DNA sequence is 94% similar. A third variant, H2A.Z, exists in yeast but has only 56% amino acid sequence homology to H2A.

**Figure 1.**
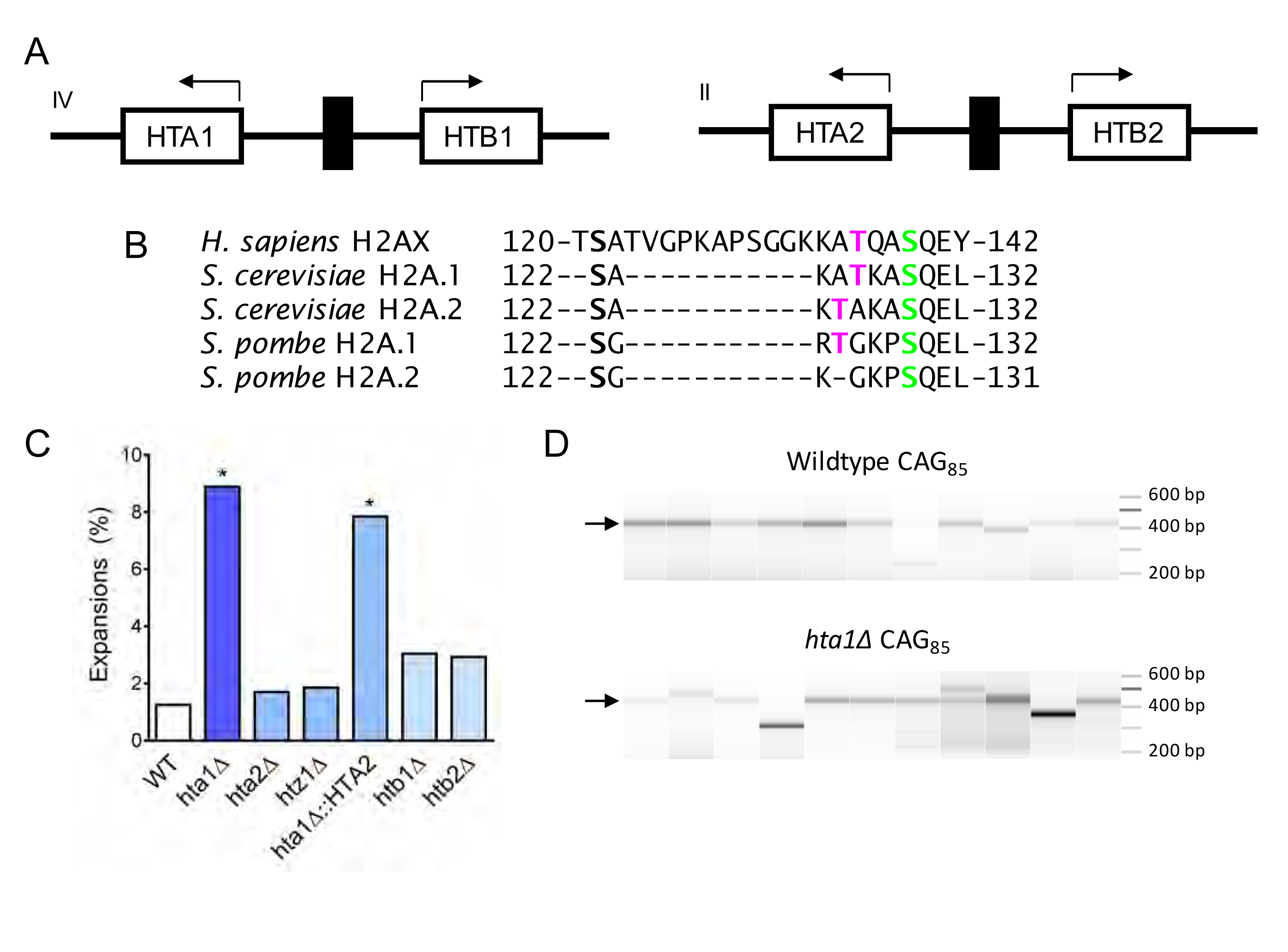
H2A.1 is required to prevent (CAG) _85_ repeat expansions. A) H2A is present in two copies in *S. cerevisiae*, paired with H2B. The gene pairs are divergently transcribed. The *HTA1-HTB1* locus is on chromosome IV, the *HTA2-HTB2* locus is on chromosome II. B) H2A/H2AX protein sequences were aligned by Clustal Omega, only the C-terminal tail residues (past the histone fold) are shown. The threonine residue at position 125/126 is present in human H2AX and conserved in *S. cerevisiae* as well as *S. pombe* H2A.1. C) CAG repeat expansion frequency was measured using a PCR-based stability assay, using primers P1 and P2, described in Figure S1A. Products were run in high-resolution Metaphor agarose or a fragment analyzer (as shown in D) and repeat length changes were evaluated. Raw instability data can be found in Table S2. Statistical deviation from wild-type was calculated by Fisher’s Exact Test, *p<0.05. D) Visualization of CAG repeat lengths using an Advanced Analytical fragment analyzer. Arrow indicates length of (CAG)_85_ PCR product.

H2A modification is a major contributor to DNA repair and may be particularly important in promoting efficient repair at unstable genomic elements. CAG trinucleotide repeats are in this category, as they can form abnormal secondary structures, such as hairpins and slip-stranded DNA (reviewed in (12-14)), and break at a higher frequency than non-repetitive DNA (15-17). Repair or replication errors within the CAG repeat can lead to instability, or a change in repeat units. Once expanded (addition of repeat units), the repeat tract is increasingly unstable and prone to further expansion in a length-dependent manner (reviewed in (13, 18)). CAG repeats are found throughout the human genome but repeat expansion beyond a threshold length of approximately 35 repeats is detrimental to the cell and can lead to human disease, including Huntington’s disease, myotonic muscular dystrophy, and several spinocerebellar ataxias (13, 19). The CAG repeat is a strong nucleosome-positioning element, shown *in vitro* by nucleosome assembly assays and visualized by electron microscopy (20, 21). The intrinsic nucleosome-positioning characteristic of the CAG repeat makes this an interesting and sensitive sequence at which to study the contribution of the chromatin environment to DNA repair. Further, the unstable nature of the repeat allows us to experimentally test the importance of chromatin and repair factors in promoting high-fidelity repair, since repair errors (errors in synthesis, alignment, processing, etc) can lead to repeat tract length changes.

Secondary structures that occur at the CAG repeat can interfere with DNA transactions, causing stalled or collapsed replication forks, gaps, nicks, and DSBs (13). Homologous recombination can be initiated to repair these lesions, but this repair itself can be a source of mutagenesis if it does not proceed with high fidelity (reviewed in (22)). Several steps during HR presumably require nucleosome repositioning or eviction, including resection, strand invasion, copying the template and D-loop extension, and resetting the chromatin structure once repair is complete. Efficient completion of each of these stages of HR is expected to be important for preventing errors that can lead to CAG repeat expansions (22).

We previously described a role for histone H4 acetylation in protecting CAG repeat stability by promoting high-fidelity homologous recombination during post-replication repair (23). Here, we explore the role of histone H2A in CAG repeat maintenance. In a primary genetic screen for CAG fragility and a secondary screen of CAG instability, deletion of histone H2A.1 was found to increase CAG repeat fragility and expansion frequency. However, deletion of the second copy of this protein, H2A.2, had no effect on repeat fragility or instability. Since histone H2A could be participating in one or more of several pathways that contribute to repeat stability, several hypotheses have been explored to explain this discrepancy. We found that H2A.1 threonine 126 (T126) is required to prevent CAG expansions and that expansions that arise in the absence of phosphorylatable T126 are dependent on Rad51, Rad52, Rad57, and the Polδ subunit Pol32. In addition, we show that H2A.1 and H2A.1-T126 are required for efficient SCR at non-repetitive DNA sequences, and are working in the same pathway as the Polδ subunit Pol32. Together, these results demonstrate that H2A.1 plays an important role in promoting efficiency and fidelity of recombination during repair. This role is distinct from H2A.2, and our results implicate Thr126 phosphorylation as important for this distinction.

## Results

### H2A histone variants contribute differentially to (CAG) _85_ repeat stability

To identify factors important for maintaining expanded CAG/CTG trinucleotide repeats, a screen was performed for factors that protect against repeat fragility using a yeast artificial chromosome (YAC) end loss assay (Figure S1A; reviewed in (24)). Initial semi-quantitative assays showed a 2-fold increase in the rate of 5-FOA-resistance (FOA^R^) in the *hta1Δ* strain compared to the wild-type for a strain containing a YAC with a (CAG)_85_ repeat tract, whereas the *hta2Δ* mutant did not deviate from wild-type (Fig S1B). Upon further analysis using multiple independent *hta1Δ* transformants, we observed a wide range of repeat fragility rates and thus were unable to statistically verify the increase over wild-type (Figure S1B; Table S1). We conclude that there is likely a mild defect in DSB repair at the CAG repeat in the absence of H2A copy 1.

We next evaluated the contribution of the H2A copies as well as the other histone H2 variants to CAG/CTG repeat stability. Histone genes were deleted and (CAG)_85_ repeat tract length changes were monitored by PCR analysis (16, 23) (Figure 1D, S1A, Table S1). Deletion of the two copies of H2A differentially affect CAG repeat stability: expansion frequency is significantly increased in the *hta1Δ* mutant (7-fold increase over wild-type, p =2.7 × 10^−5^) while expansion frequency in the *hta2Δ* mutant is not significantly changed from wild-type levels (1.3-fold increase over wild-type, p = 0.71) (Figure 1C). Contractions were increased 2.6-fold in the *hta1Δ* but not the *hta2Δ* mutant (Table S1). Thus, H2A.1 is required to suppress CAG instability while H2A.2 is not.

The H2A.Z variant is encoded by the gene *HTZ1* in yeast. Since H2A.Z is deposited at a DSB during repair (25), we tested whether it is required to maintain CAG repeat stability. Although the *htz1Δ* mutant is sensitive to DNA damaging agents (26, 27), deleting the H2A.Z gene did not affect repeat expansion frequency, and is therefore dispensible for CAG repeat maintenance (Figure 1C) (23). Likewise, deletion of either H2B gene had no significant effect on repeat expansion frequency (Figure 1C). Therefore, of the histone H2 proteins, only H2A.1 plays a significant role in preventing CAG repeat expansions.

### H2A.1 sequence, not histone levels, confers specificity to its role in preventing CAG repeat expansions

Histone H2A is encoded by two nearly identical gene copies in *S. cerevisiae,* each paired with a copy of H2B and differentially regulated (28) (Figure 1A). While the *HTA1-HTB1* (TRT1 locus) gene pair can be upregulated in the absence of *HTA2* to maintain the correct stoichiometric ratio of H2A in the cell, *HTA2* is transcribed at a constant rate (29). As a result, the H2A pools will be normal in an *hta2Δ* mutant, whereas an *hta1Δ* mutant may have a global decrease in H2A. A second pathway of gene dosage compensation has been documented by which the *HTA2-HTB2* gene pair amplifies to form a minichromosome (that also contains the *HHT1-HHF1* (H3-H4) gene pair) in the absence of *HTA1-HTB1* (30). To distinguish if repeat stability is mediated by the H2A protein sequence or histone levels, the *HTA2* sequence was placed under the control of the *HTA1* promoter, replacing the *HTA1* gene (*hta1Δ::HTA2*). In this strain, the H2A.2 protein will be expressed at the same level and timing as H2A.1 in a wild-type cell but the H2A.1 protein will not be present. Equal expression of H2A proteins was confirmed by Western blot (Fig. S2A). If H2A expression level or timing is the major contributor to repair of the CAG repeat, repeat maintenance will be at wild-type levels when H2A.2 is expressed from the H2A.1 promoter. However, we observed that CAG repeat expansions remained significantly increased from the wild-type in the *hta1Δ::HTA2* strain (Figure 1C), though a partial suppression of contractions was observed (Table S1). Therefore, H2A.2 cannot compensate for H2A.1 in preventing repeat expansions, even when expressed at H2A.1 levels under control of the *HTA1* promoter. We conclude that the sequence of H2A.1, not histone levels or subtleties in expression timing, is required to prevent CAG repeat expansions.

The H2A.1 and H2A.2 protein sequences are identical except for a direct threonine/alanine (T/A) switch in the C-terminal tail (Figure 1B). Despite this high sequence identity, the histone subtypes have been documented before to play different functions during the *S. cerevisiae* life cycle. The absence of gene product from *HTA1* and *HTB1* (*TRT1Δ*) leads to a constitutive heat shock response after exposure to to high temperature, and this phenotype was not rescued with an additional copy of *HTA2* and *HTB2* (TRT2) (28). After heat shock in human cells, there is an increase in fork arrest or slowing in S-phase that is marked by γH2AX, while in G1 and G2 there is a measurable increase in DSBs (31). Taken with our results demonstrating differential roles for the H2A copies in maintaining CAG repeats, this further supports that H2A.1 is specifically important for repair or recovery from DNA damage.

### Nucleosome positioning at the CAG repeat is maintained in H2A mutants

Since the H2A.2 gene is not upregulated in an H2A.1 deletion mutant, we hypothesized that an *hta1Δ* mutant may cause histone depletion or disruption of the chromatin structure at the CAG repeat tract, leading to increased instability. Although overall bulk chromatin structure was not altered in an *hta1Δhtb1Δ* strain, some areas of the genome were more sensitive to micrococcal nuclease (MNase) digestion in the absence of H2A.1 (32). The CAG repeat is a strong nucleosome positioning element and thus could be more sensitive to H2A.1 depletion than other regions of the genome. To visualize nucleosome positioning at the CAG repeat, we used indirect end-labeling with a probe upstream of the CAG repeat (Figure S1A, red line) and measured sensitivity of the chromatin to MNase digestion. The result shows a series of discrete, protected fragments (approximately five nucleosomes), indicating several positioned nucleosomes within and flanking the CAG repeat (Figure 2A). The pattern is consistent with a canonical ∼165 bp spacing between nucleosomes, and this does not appear to be significantly altered in *hta1Δ* cells compared to the wild-type (Figure 2A).

**Figure 2.**
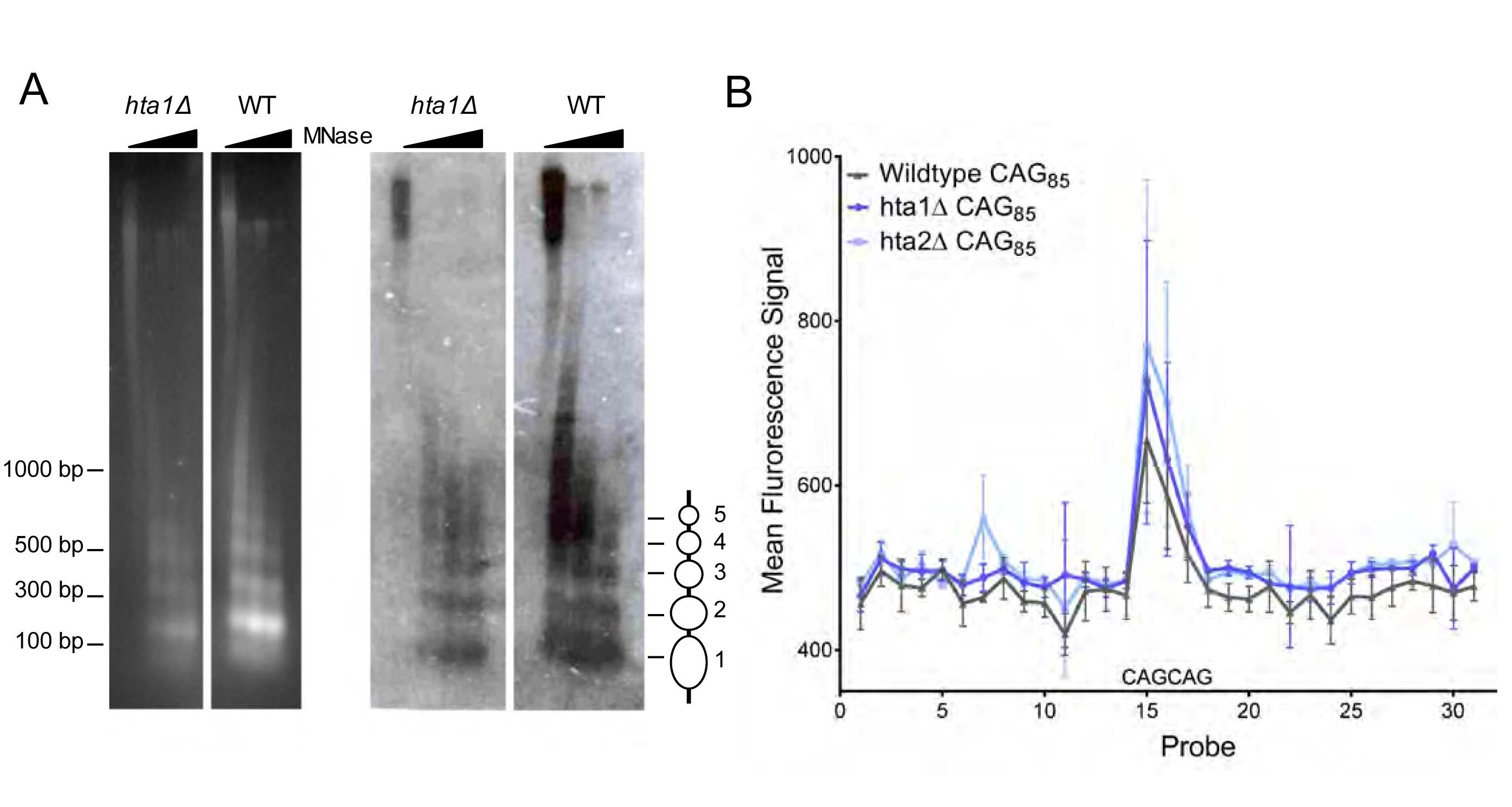
Nucleosome positioning at a (CAG) _85_ repeat is not altered in the absence of H2A.1 or H2A.2. A) Indirect end-labeling of nucleosomal DNA upstream of the CAG repeat. MNase (0, 0.25, 2.5, and 7.5 units) digested DNA was run in 1.5% agarose with ethidium bromide (left) and Southern blotted (right) using a probe ∼100 bp proximal to the CAG repeat (red line Figure S1A). Ovals represent nucleosome positions. B) Illumina array mapping of nucleosome protection at the CAG repeat. Mononucleosomal DNA from strains containing the (CAG)_85_ repeats was hybridized to a custom array of 30-mer probes spanning 425 bp upstream of the repeat to 436 bp downstream of the repeat in YAC CF1. Probes 14-16 contain CAG repeats; probe 15 is composed purely of CAG repeats (probe sequences in Table S3).

To generate a high-resolution nucleosome map of the regions flanking the CAG repeat, we used a custom Illumina BeadArray containing probes spanning 425 bp upstream to 438 bp downstream of the CAG repeat tract on the YAC, including two CAG repeat containing probes and one pure CAG repeat probe, in non-overlapping 30-mers (Table S3). We hybridized MNase digested mononucleosomes of *hta1Δ, hta2Δ*, and wild-type (CAG)_85_ cells to the array to measure nucleosome protection at the CAG repeat. The CAG repeat-containing probes produced a peak in intensity compared to the flanking non-CAG repeat-containing probes, indicating strong nucleosomal protection at the CAG repeat (Figure 2B). This result confirms previous *in vitro* data that the CAG repeat is a strong nucleosome binding sequence (21, 33, 34) and shows that it positions a nucleosome *in vivo* on a yeast chromosome. In addition, the protection was not reduced in the *hta1Δ* and *hta2Δ* mutants (Figure 2B). Thus, both methods used show that there is a positioned nucleosome at the CAG tract that is not altered in the *hta1Δ* background. We conclude that nucleosome positioning is not the major contributing factor to CAG instability in the *hta1Δ* mutant.

### A phosphorylatable Thr126 residue in the H2A.1 C-terminal tail is required for (CAG) _85_ repeat maintenance

H2A.1 and H2A.2 vary in amino acid sequence only by the position of threonine in the C-terminal tail, which occurs at either position 126 in H2A.1 or 125 in H2A.2 (Figure 1B). To address the role of specific residues in the H2A C-terminal tail in preventing CAG expansions, we used strains in which the genomic copies of H2A and H2B are deleted and H2A point mutants are expressed from a plasmid that also contains a wild-type copy of *HTB1* (35). In this system, we observed a slightly elevated frequency of (CAG)_85_ expansions in *HTA1-WT* plasmid strain: 2% versus 1.3% (Table S1). This is likely due to the presence of a plasmid rather than histone dosage, as a similar phenomenon has been observed with other plasmid-containing strains (23). When the threonine at position 126 was rendered non-modifiable by mutation to alanine, expansions were significantly increased over the *HTA1*-plasmid wild-type (6.1%; 3.1-fold over wt, p = 0.04) (Figure 3A). Integration of the *T126A* point mutation at the endogenous *HTA1* locus resulted in a similar expansion frequency when H2A.2 is not present (3.7-fold over wt, Table S1). CAG contractions were relatively unaffected, therefore this residue primarily protects against expansions. Additionally, when the threonine at position 126 was mutated to glutamic acid to mimic constitutive phosphorylation, expansions were significantly increased over wild-type (4.9%, 3.7-fold over WT, p=0.01) (Table S2). This suggests not only that a phosphorylatable threonine is necessary for stability of the CAG repeat tract, but that dynamic phosphophylation of H2A.1-T126 is important. Taken together with the specific role for H2A.1, this result suggests that the position of the threonine within the tail affects the efficiency with which the H2A copies contribute to DNA replication or repair.

**Figure 3.**
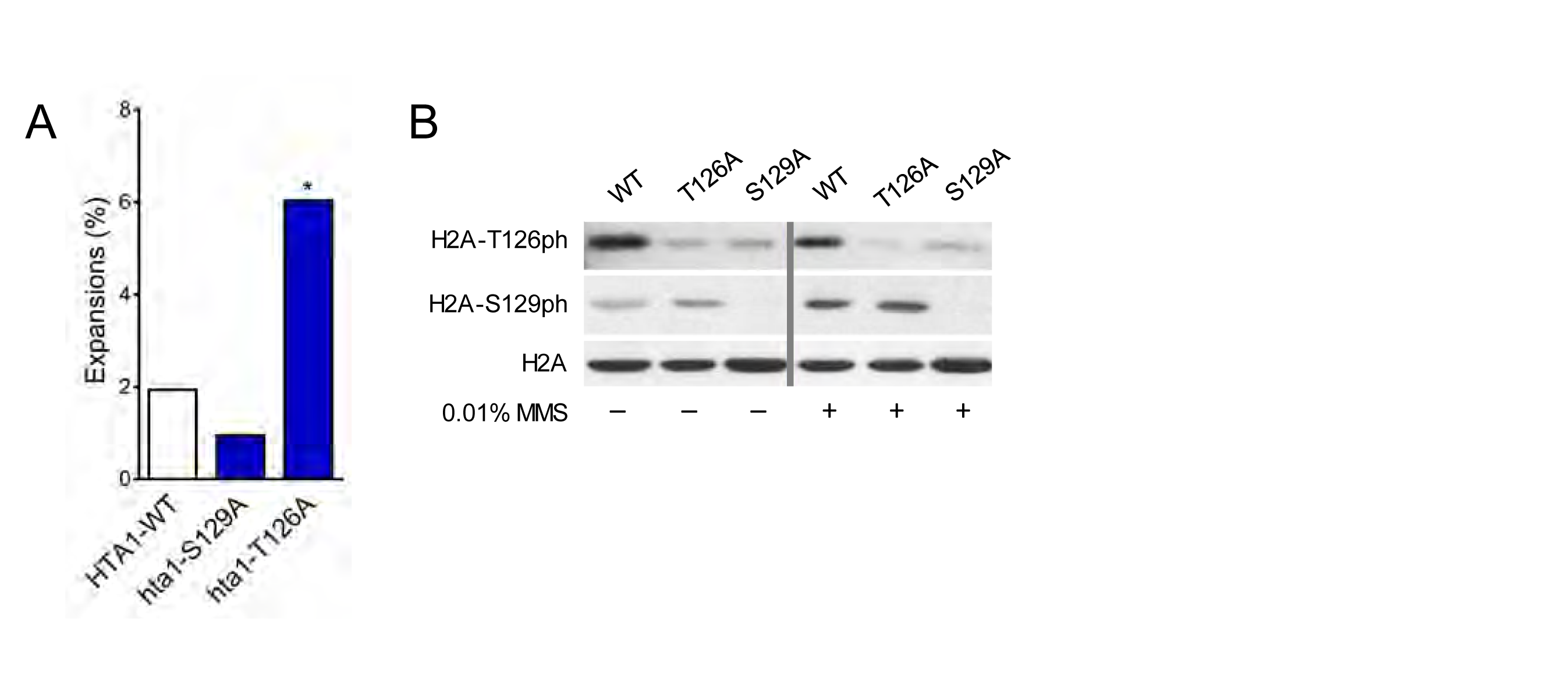
H2A.1-T126 is required to prevent (CAG) _85_ repeat expansions and is phosphorylated ***in vivo***. A) CAG repeat length changes were evaluated in strains with H2A.1 C-terminal tail point mutations. Strains contain only one copy of H2A.1, either wild-type (WT) or with the indicated point mutation, expressed from a plasmid, and no copy of H2A.2. Statistical deviation from the strain containing a plasmid with wild-type HTA1 was calculated by Fisher’s Exact Test, *p<0.05. B) Phosphorylation of H2A-T126 and S129 was evaluated by Western blot in the H2A wild-type and point mutant strains. Antibodies specific to the phospho residues as indicated detected phosphorylation in untreated cells (left panel) or cells that had been treated with 0.01% MMS (right panel).

We also tested the importance of H2A-S129 phosphorylation in repeat maintenance. Despite H2A-S129ph being preferentially detected at a (CAG)_155_ tract in this same location by ChIP (23), the ability to phosphorylate H2A-S129 did not significantly affect CAG repeat tract stability, as neither contraction nor expansion frequencies were significantly altered from wild-type in the *hta1-S129A* plasmid point mutant or in a genomic integration of the *S129A* mutation (Figure 3A; Table S2).

### **H2A-T126 is phosphorylated *in vivo***

In order to evaluate *in vivo* phosphorylation of the H2A C-terminal tail residues, we used a custom antibody raised against a phosphopeptide corresponding to the H2A C-terminal tail phosphorylated at T126. The antibody specifically recognizes H2A C-terminal tail peptides phosphorylated at T126, but not unphosphorylated peptides or peptides phosphorylated at S122 or S129 (Fig. S3). This antibody is also fairly specific *in vivo*, as there is low background in the *hta1-T126A* mutant (Figure 3B, second lane) and it detects only H2A.1 but not H2A.2 (Figure S2B). We found that H2A-Thr126 is phosphorylated in wild-type cells even in the absence of DNA damaging agents, in agreement with previous results using a different antibody preparation that is no longer available (36, 37). Interestingly, the H2A-T126ph band is less intense in the *hta1-S129A* mutant, demonstrating that the *H2A-S129A* mutation impairs binding of the T126ph antibody (Figure 3B). This result indicates that the T126ph antibody may need an accessible serine at position 129 (either phosphorylated or not) for full binding. Importantly, recognition of H2A-S129ph is not impaired by the *hta1-T126A* mutation. Therefore, H2A-S129ph may occur unimpeded when T126 is unphosphorylated and deficient γH2A formation cannot explain the phenotype of the *hta1-T126A* mutant.

We next sought to determine if phosphorylation of H2A-T126 changes upon exposure to MMS, a DNA base alkylating agent that causes abasic sites that can be converted into single and double strand breaks. Previously, the *hta1-T126A* mutant was shown to have a mild survival defect when spotted on 0.02% MMS and T126 phosphorylation was slightly increased in the presence of MMS (37). In the presence of 0.01% MMS, H2A-S129ph is increased, as expected, but H2A-T126ph is unchanged and is therefore not induced by the type of damage caused by low levels of MMS (Figure 3B). Treatment with a higher level of MMS (0.03%), 0.2 M hydroxyurea (HU), or HU+MMS (0.2M, 0.03%) also did not alter H2A-T126 phosphorylation levels in our strains (Figure S2D). We conclude that H2A-T126 phosphorylation is not a damage-induced modification. Consistent with this conclusion, T126ph levels remained unchanged in the absence of the major DNA-damage inducible kinases (*mec1Δsml1Δ, tel1Δ, rad9Δ, rad53-21, chk1Δ*, and *dun1Δ* mutants; Figure S2C).

### H2A.1 promotes fidelity of homologous recombination (HR) events

To determine how H2A.1 and H2A-T126 might be contributing to fidelity of CAG repeat repair and preventing expansions, we assessed repeat stability in the absence of DNA repair pathways. If expansions in the *hta1Δ* mutant are arising through a low-fidelity repair event, repeat expansions will be abolished in the absence of the relevant repair pathway.

We first tested if expansions in the absence of H2A.1 are arising through NHEJ by deleting Lif1, a DNA ligase IV subunit, in the *hta1Δ* and *hta1-T126A* mutants. Expansion frequency remained elevated in the *hta1Δlif1Δ* and *hta1-T126A lif1Δ* double mutants (Figure 4A, 4B); therefore, instability in the absence of H2A.1 is not arising through low-fidelity NHEJ. Consistently, Moore et al found no NHEJ defects in a plasmid end-joining assay in an *hta1-T126A* mutant compared to a strain with wild-type H2A (37). We next surveyed homology-dependent repair pathways. Chromatin modifications have recently been shown to influence repair fidelity of post-replication repair (23), and so we assayed CAG repeat stability in the *hta1Δrad5Δ* and *hta1-T126A rad5Δ* double mutants. Although expansions are somewhat suppressed in the *hta1Δrad5Δ* double mutant, H2A.1 does not appear to be significantly contributing to the fidelity of post-replication repair (Figure 4A). Likewise, expansions remain elevated in the *hta1-T126A rad5Δ* double mutant (Figure 4B). To determine if CAG expansions in the absence of H2A.1 are arising through general HR, we measured stability of the CAG repeat in the absence of two key HR proteins: Rad52 and Rad51. Rad52 binds RPA-coated ssDNA and loads Rad51 to form the Rad51 filament that will allow homology search during HR repair. Expansions in the *hta1Δrad52Δ* double mutant are significantly reduced 2.9-fold from the *hta1Δ* single mutant, and a similar suppression is seen in the *hta1-T126A rad52Δ* double mutant, though not to the level of statistical significance (Figure 4A, 4B). Similarly, expansion frequencies in the *hta1Δrad51Δ* and *hta1-T126A rad51Δ* double mutants are suppressed from the single mutant expansion frequencies (Figure 4A, B). Corroborating these results, expansions were suppressed almost to wild-type levels in the *hta1Δrad57Δ* mutant (Figure 4A) and were suppressed approximately 3-fold in the *hta1-T126A rad57Δ* mutant. Together, these results indicate that expansions in the absence of H2A.1 are arising through Rad51-and Rad52-dependent HR events and implicate T126 phosphorylation in regulating this role. Since Rad57 is especially required for SCR (38), the dependency on Rad57 is consistent with expansions arising during SCR, a pathway previously implicated in causing CAG instability (39, 40).

**Figure 4.**
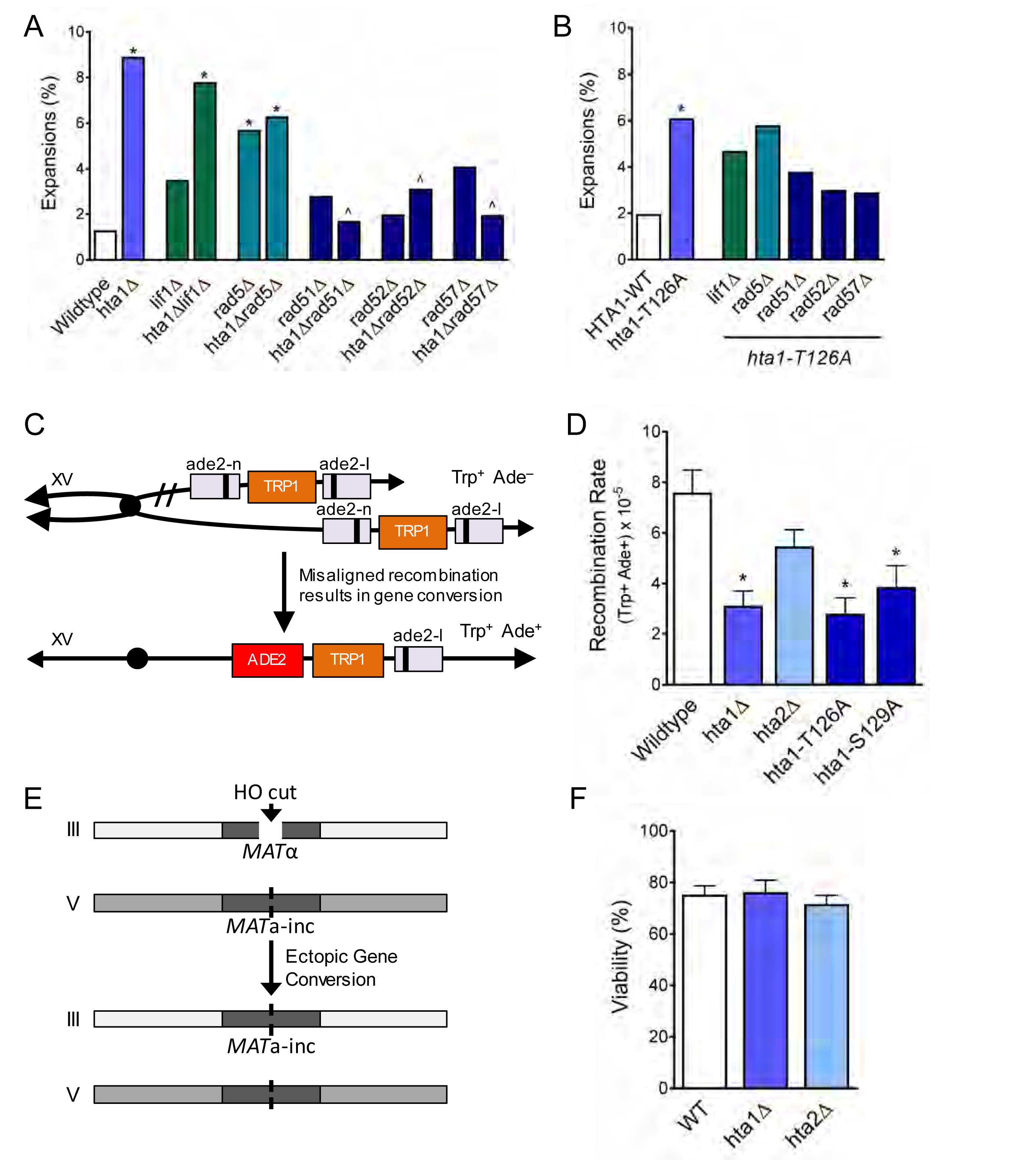
H2A.1 promotes fidelity of HR to prevent (CAG) _85_ expansions and is required for efficient SCR. A) Changes in CAG repeat length were assessed as in Figure 1C. Statistical deviation from wild-type was calculated by Fisher’s Exact Test, *p<0.05. B) Repair proteins were deleted in the strain expressing *hta1*-*T126A* from the plasmid; no copy of H2A.2 is present in these cells. Changes in CAG repeat length were assessed as in (A). Statistical deviation from the plasmid *HTA1* wild-type were calculated by Fisher’s Exact Test, *p<0.05. C) Misaligned recombination during SCR can be measured by gene conversion from Trp^+^ Ade^−^ to Trp^+^ Ade^+^. Only gene conversion of the lower chromatid is shown (23, 38). D) Rates of spontaneous unequal SCR. For these experiments, the *hta1-T126A* and *hta1-S129A* mutations were integrated at the genomic locus, replacing the wild-type copy of *HTA1*; *HTA2* remains intact in these strains. SCR rates for the H2A.1 point mutants with HTA2 deleted appear in Table S4. Statistical deviations as indicated were calculated using a Student’s t-test, *p<0.05 to wild-type. E) Ectopic gene conversion at a DSB can be measured by viability in the presence of galactose-induced HO. The *MAT*α locus on chromosome 3 contains the HO cut site, whereas the *MAT***a**-*inc* locus on chromosome 5 has no HO cut site. Conversion to *MAT***a**-*inc* on chromosome 3 by ectopic gene conversion eliminates the HO target site and cells can survive in the presence of galactose (42). F) Rates of ectopic gene conversion. Statistical deviation from wild-type was tested by a Student’s t-test.

### H2A.1 and H2A-Thr126 promote efficient SCR

To further evaluate the role of H2A.1 in homology-mediated repair events, we assayed the H2A mutants for their ability to modulate SCR using a genetic assay that measures rates of spontaneous unequal SCR as an estimate of overall SCR levels (38) (Figure 4C). SCR is not suppressed from wild-type in the *hta2Δ* mutant, but is significantly suppressed in the *hta1Δ* mutant (2.4-fold suppression from wild-type, p = 7.9 × 10^−3^; Figure 4D). Thus, H2A.1 is required for efficient SCR while H2A.2 is not, mirroring the differential role of the two H2A copies in CAG repeat maintenance. Similarly, SCR levels are decreased 2.7-fold from wild-type in the *hta1-T126A* mutant (p = 4.9 × 10^−3^) (Figure 4D). SCR is also suppressed, though more mildly, in the *hta1-S129A* mutant (by 1.9-fold from wild-type; p = 0.03; Figure 4D), in agreement with a previous report that found a mild defect in SCR during repair of a DSB in an *hta1-S129A* mutant (41). Rates of SCR in the *hta1-T126A* and *hta1-S129A* mutants are also reduced in the *hta2Δ* background (Table S4), indicating that this reduction does not depend on the presence or absence of H2A.2. These results demonstrate that H2A.1 and H2A-T126 are required for efficient spontaneous SCR. This supports our conclusion that H2A.1 and H2A.1-T126 are required for proper recombination at the CAG repeat tract to prevent repeat expansions, and extends this finding to non-repetitive DNA sequences.

Given the role for H2A.1 in promoting spontaneous SCR, a process that is likely to be predominantly initated by single-stranded gaps at replication forks (38), we were curious if H2A.1 also promotes recombination at DSBs. To measure this, we employed an assay by which an HO-induced DSB at the *MAT*α locus on chromosome 3 can be repaired via ectopic gene conversion from a *MAT***a**-*inc* donor on chromosome 5 (42, 43) (Figure 4E). The *MAT***a**- *inc* allele cannot be cut by the HO-endonuclease, and therefore rates of survival in the presence of galactose-inducible HO can be used as a measure of interchromosomal recombination frequency at a DSB. We found no difference in viability between *hta1Δ, hta2Δ*, and wild-type (Figure 4F; Table S5). Therefore, there is no significant H2A.1-specific role in promoting interchromosomal HR at DSBs, and this strengthens our conclusion that H2A.1 is important for recombination between sister chromatids.

### H2A.1 functions with Pol32 during SCR and promotes efficient break-induced replication (BIR)

We were interested in which step of SCR H2A.1 was involved. Since CAG expansions could occur during DNA synthesis, we tested the role of the Polδ subunit Pol32, which is required for Polδ processivity. Interestingly, SCR is significantly suppressed from the wild-type in the *pol32δ* mutant. This establishes that the Pol32 subunit of PolΔ is required for D-loop extension during the short tract recombination measured by this assay (less than 1 kb). Notably, SCR suppression in the *pol32Δ* mutant is epistatic to the absence of H2A.1 and a phosphorylatable H2A.1-T126 residue, as the SCR rate is not further diminished in the *hta1Δpol32Δ* or *hta1-T126A pol32δ* double mutants (Figure 5A). This suggests that H2A.1 and Thr126 phosphorylation may be important in facilitating D-loop extension by Polδ during recombination.

**Figure 5.**
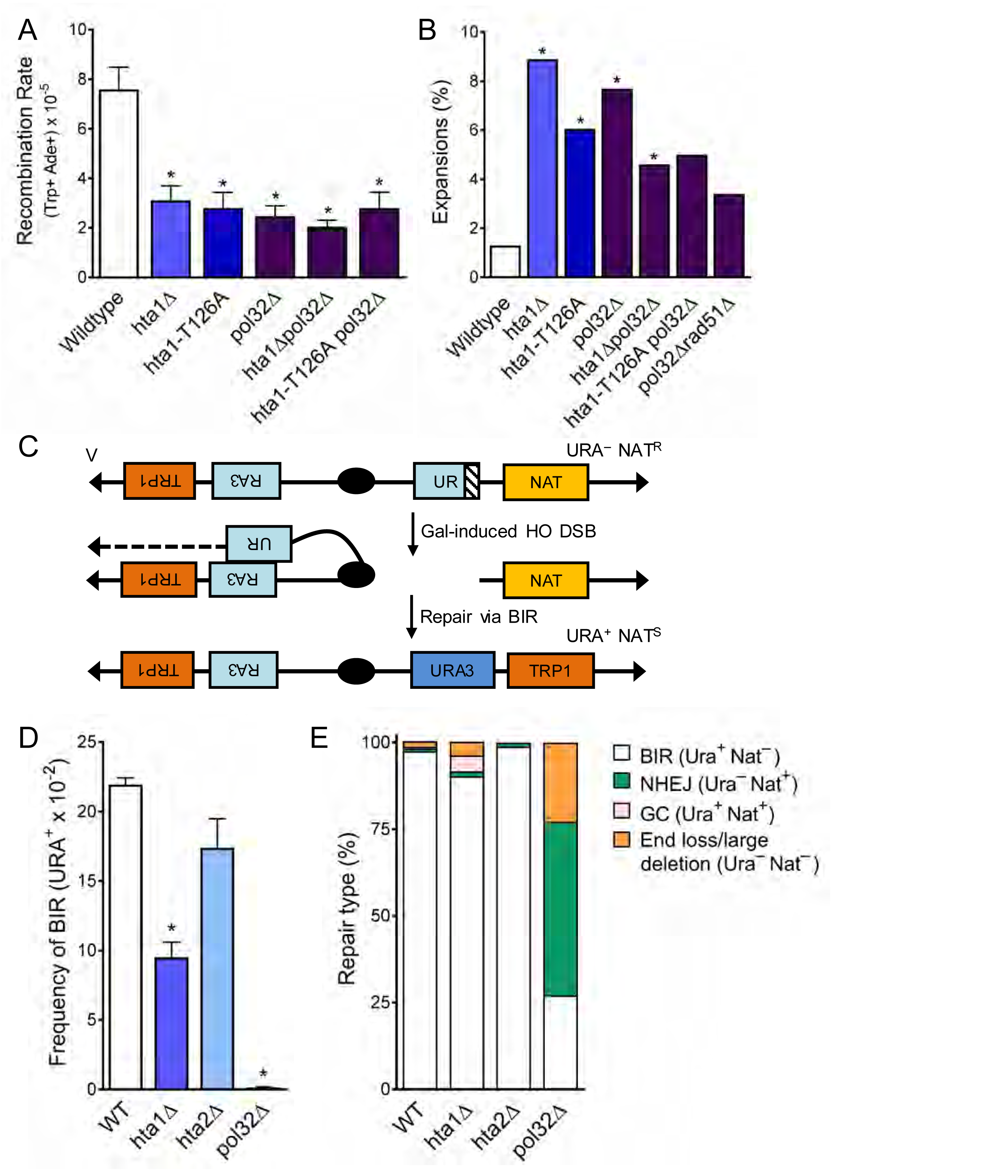
H2A.1 and Pol32 work in the same pathway during SCR, and H2A.1 is required for efficient BIR. A) Rates of spontaneous unequal SCR, assessed as in Figure 4D. Statistical deviation from wild-type was calculated using a Student’s t-test, *p<0.05. B) Changes in CAG repeat length were assessed as in Figure 1C. Statistical deviation from wild-type was calculated by Fisher’s Exact Test, *p<0.05. C) Assay to measure BIR at an HO-induced DSB. The first 400bp of the URA3 gene (UR; right arm) is upstream of an HO cut site (stripes). Homology driven repair of the HO induced DSB can occur with the remaining 404 bp of the URA3 gene (A3; left arm) and if completed via BIR renders the cell URA^+^ and NAT^S^ (47). D) Rates of BIR during repair of an HO-induced DSB. Statistical deviation from wildtype was calculated using a Student’s t-test, *p<0.05, **p<0.01. E) The type of repair induced by the HO DSB was evaluated by individually scoring all colonies for growth on YC-URA and YEPD+Nat. Colonies that were URA^+^ NAT^S^ are were repaired via BIR. Other types of possible repair events included gene conversion (URA^+^ NAT^R^), non-homologous end-joining (URA^-^ NAT^R^), and *de novo* telomere addition (URA^-^ NAT^S^).

We next tested the role of Pol32 in CAG expansions. In the *pol32δ* single mutant, CAG repeat expansions are significantly increased over wild-type (5.8-fold over wild-type, p =2.0 × 10^−3^), indicating that processive DNA synthesis by Polδ is required to prevent repeat expansions (Figure 5B). We tested whether instability in the absence of Pol32 was due to its role in normal replication or recombination-associated DNA synthesis by deleting Rad51 in a *pol32*Δ mutant (Figure 5B). Expansions in the *pol32Δ* mutant were suppressed in the absence of Rad51 (to 3.4%, 2.3-fold decrease from *pol32*Δ, p = 0.17), suggesting that expansions are at least in part due to polymerase slippage during D-loop synthesis. The increase in expansion frequency in the *pol32Δ* mutant is similar to that in the *hta1Δ* and *hta1-T126A* mutants. However, CAG expansion frequency drops below the level of each single mutant in the *hta1Δpol32Δ* and *hta1-T126A pol32Δ* mutants (Figure 5B), suggesting that Pol32-dependent synthesis is responsible for some of the expansions occurring in the *hta1Δ* background. Though the suppression is not statistically significant (for example p=0.12 for *hta1Δpol32Δ* compared to *hta1Δ*), it is markedly reduced from the additive levels expected if there was no interaction between the pathways. Since the expansions in *hta1Δ* and *hta1-T126A* are suppressed in the absence of Rad51, Rad52, Rad57, and Pol32 (Figure 4A, 5B), they are likely arising downstream of synapsis and initiation of DNA synthesis during recombination. Taken together, these results suggest that the CAG expansions observed in the absence of H2A.1 occur during a Pol32-dependent recombination process. For example, H2A.1 could promote efficient D-loop extension during replication of the donor strand, thereby preventing opportunity for DNA secondary structure formation.

Pol32 is known to be especially important for BIR, which involves extended D-loop synthesis that can proceed for many kilobases (44-46) (see also Figure 5D). Considering the importance of Pol32 in preventing CAG expansions and the evidence that H2A.1 and Pol32 are functioning in the same pathway during SCR and prevention of CAG expansions, we wondered whether H2A.1 might also have a role in BIR. To directly test the role of H2A.1 in BIR, we used a system in which a DSB induced by the HO endonuclease can result in a non-reciprocal translocation when repair proceeds via BIR (47) (Figure 5C). BIR frequency is suppressed 2.3-fold from wild-type in the *hta1Δ* mutant, but remains at wild-type levels in the *hta2Δ* mutant (Figure 5D; Table S6). The repair outcome profile in the *hta1Δ* mutant is also altered from wild-type: a greater proportion of cells either undergo gene conversion to become URA^+^ Nat^+^ or lose both the URA gene and the NAT^R^ marker, which can occur due to *de novo* telomere addition after DSB induction or inaccurate end joining that inactivates the NAT gene (Anand and Haber, personal communication) (Figure 5D). Therefore, deletion of H2A.1 in the BIR system results in decreased BIR and alterations in repair type frequencies, demonstrating that H2A.1 plays a role in facilitating efficient BIR. Together with the SCR data, this suggests that expansions in the *hta1Δ* mutant may be due to defective D-loop extension, since this step occurs during both SCR and BIR.

## Discussion

Stemming from an initial observation in a genetic screen that CAG repeat fragility and instability were elevated in an *hta1Δ* mutant but not an *hta2Δ* mutant, we demonstrated that H2A.1 and H2A.2 differentially contribute to homology-mediated repair of CAG repeats. Significantly, we found that repeat expansions in the *hta1Δ* and *hta1-T126A* mutants depend on Rad51, Rad52, and Rad57, indicating that H2A.1 and T126 are required for high-fidelity HR repair (Figure 4A). Further, H2A.1 promotes efficient SCR (Figure 4D) and BIR (Figure 5D, E). Nucleosome positioning at the CAG repeat cannot account for the different roles of H2A.1 and H2A.2, as the CAG repeat is highly nucleosome protected in both the *hta1Δ* and *hta2Δ* mutants (Figure 2A, B), and the elevated expansion frequency in the absence of H2A.1 is not rescued when H2A.2 is expressed at H2A.1 levels (Figure 1C). Therefore, our data support that H2A.1 may be more efficient at promoting high-fidelity HR repair than H2A.2 because of the protein sequence; specifically, that the position of the phosphorylatable threonine in the H2A.1 C-terminal tail is more advantageous to repair than in H2A.2. Both copies of yeast H2A contain the SQEL motif in the C-terminal tail, and therefore both copies are considered homologs of mammalian H2AX. A compelling implication of our result here is that yeast H2A.1 is in fact the closer homolog of mammalian H2AX, as it plays a greater HR-specific role than H2A.2. Like H2A.1, mammalian H2AX also contains a phophorylatable threonine two residues before serine 139. Also similar to H2AX, H2A.1 is initially a smaller proportion of the total H2A pool (29, 48). Consequently, H2A.1 and H2A.2 may be more akin to histone variants than histone copies.

### A role for H2A.1 T126 in promoting high-fidelity recombination-mediated repair

H2A-Thr126 has previously been shown to be phosphorylated *in vivo* but its phosphorylation state after DNA damage and its overall contribution to break repair was unclear (35-37, 49). Using a naturally unstable expanded CAG repeat tract, we have shown that phophorylation of T126 is important for maintaining CAG repeats and plays a role in promoting efficient SCR. Our results indicate that overall levels of HR/SCR are suppressed in the absence of phosphorylatable H2A.1-T126, and that the recombination that does take place proceeds with low fidelity, leading to repeat expansions. H2A.1-T126ph is fairly abundant in the cell even in the absence of DNA damage, and bulk levels of T126ph were not altered in response to the DNA damaging agents tested here (Figure 3B, S2D). This suggests that H2A.1-T126ph may be a pre-existing histone modification required for repair, rather a damage-inducible modification. H2A.1-T126ph could influence one or more steps of recombination at the CAG repeat, explored below.

The fact that expansions, an addition of bases, occur in *hta1Δ* and *hta1-T126A* mutants is most supportive of a role for H2A.1 in promoting a synthesis step of DNA repair. Also, these expansions are likely arising downstream of synapsis and D-loop assembly since they were suppressed in the absence of Rad51, Rad52, and Rad57 (Figure 4A). Further, our data placing H2A.1 and T126 in the same pathway as Pol32 in the unequal SCR assay supports a role during D-loop extension. Therefore, we conclude that H2A.1 is most likely promoting efficient D-loop extension during replication of the donor strand. A second possibility is that H2A.1-T126 is required to promote sister chromatid cohesion, preventing out-of-register alignment during recombination with the sister chromatid and repeat number changes. However, we do not favor this model as cohesin (Mcd1 and Smc1) and cohesin loader (Scc2) subunits colocalize with γH2A by ChIP at a DSB but this recruitment requires H2A-S129ph (50), and the *hta1-S129A* mutant did not show an increased repeat expansion frequency. Second, neither the *scc1-73* cohesin mutant nor the *scc2-4* cohesin loading mutant displayed an increase in CAG expansion frequency (51).

How could a histone modification promote error-free repair? One possibility is that H2A.1-T126ph interacts with one or more chromatin remodelers to facilitate nucleosome movement or displacement. Interestingly, in G2/M when a sister chromatid is available for recombination, γH2A is dispensible for recruitment of several chromatin modifiers, including Ino80 and subunits of RSC, NuA4, Rpd3, SWI/SNF, and SWR-C (52). This leaves the intriguing possibility that the H2A.1-T126 residue is important for recruiting (or retaining) chromatin remodelers to the CAG repeat during recombinational repair. H2A.1-T126ph-mediated recruitment of chromatin remodelers could be important for opening the chromatin structure to allow access by repair factors, to remodel chromatin on the donor strand to facilitate synapsis or D-loop extension, or for resetting the chromatin structure to promote repair resolution. The *hta1-T126A* mutant displays a defect in telomere positioning effect (TPE) (36), consistent with a role in reestablishing chromatin structure after repair. We previously established that RSC and NuA4 are required to promote high-fidelity repair of the CAG repeat, and showed that RSC subunits are recruited to the repeat during S-phase. We concluded that these factors are promoting post-replication repair events (23), whereas H2A.1-T126 does not significantly contribute to Rad5-dependent repair (Figure 4A). However, it is possible that RSC and NuA4 also have a more general role in any D-loop mediated repair process. Another interesting candidate is INO80, which contains two subunits that mediate binding to phospho-histones: Arp4 and Nhp10 (53, 54)). Recruitment of any of these proteins to the CAG repeat during repair could facilitate chromatin remodeling to promote efficient D-loop progression and high-fidelity repair.

A second, but not mutually exclusive, possibility is that H2A.1 S129 and T126 residues are involved in a crosstalk-mediated signaling pathway that contributes to DNA repair. Although H2A.1-T126ph is not induced by DNA damage (Figure 3B; S2), this modification is impeded by the presence of alanine at position 129. Replacement of serine to alanine at position 129 may disrupt the docking site by the kinase of T126 on the histone tail. Of note, this is not a two-way circuit, as H2A-S129ph does not require a phosphorylatable T126 residue. It is also possible that proteins binding to S129ph could occlude the T126ph site from modification or protein docking. For example, competition between proteins that bind to one or both phosphorylated residues could facilitate removal of S129ph-bound proteins and checkpoint dampening. This idea is supported by the similar increase in expansions in the T126A and T126E mutants, indicating that dephosphylation of T126 is important for repair fidelity.

### A model for H2A.1 in HR-mediated repair of CAG repeats

Hairpins formed by CAG or CTG repeats interfere with replication and induce joint molecules between sister chromatids, which have been visualized by two-dimensional gel electrophoresis (39, 40). We propose a model in which gaps caused by replication bypass of the CAG repeat are repaired via SCR, and that H2A.1-T126ph ensures efficient remodeling and progression of Pol32-mediated D-loop extension, leading to high-fidelity repair (Figure 6A). In the absence of H2A.1, suboptimal signaling from the H2A.2-T125 residue may inefficiently recruit or retain chromatin remodelers or other DNA repair proteins, impeding D-loop extension due to a chromatin state that is not permissive to extension and copying (Figure 6B). Inefficient progression of the recombination intermediate could lead to Polδ stalling or transient dissociation of the 3’ end of the invading strand, which would allow opportunity for CAG repeat secondary structure formation and slippage during synthesis, leading to repeat length changes in the repaired DNA strand. Alternatively, chromatin remodelers or repair factors recruited by the H2A.1-T126 modification may be required to reset the chromatin structure after repair. The permissive chromatin state could allow multiple, aberrant invasion events that would increase the opportunity for misalignments during D-loop initiation or elongation, resulting in repeat instability.

**Figure 6.**
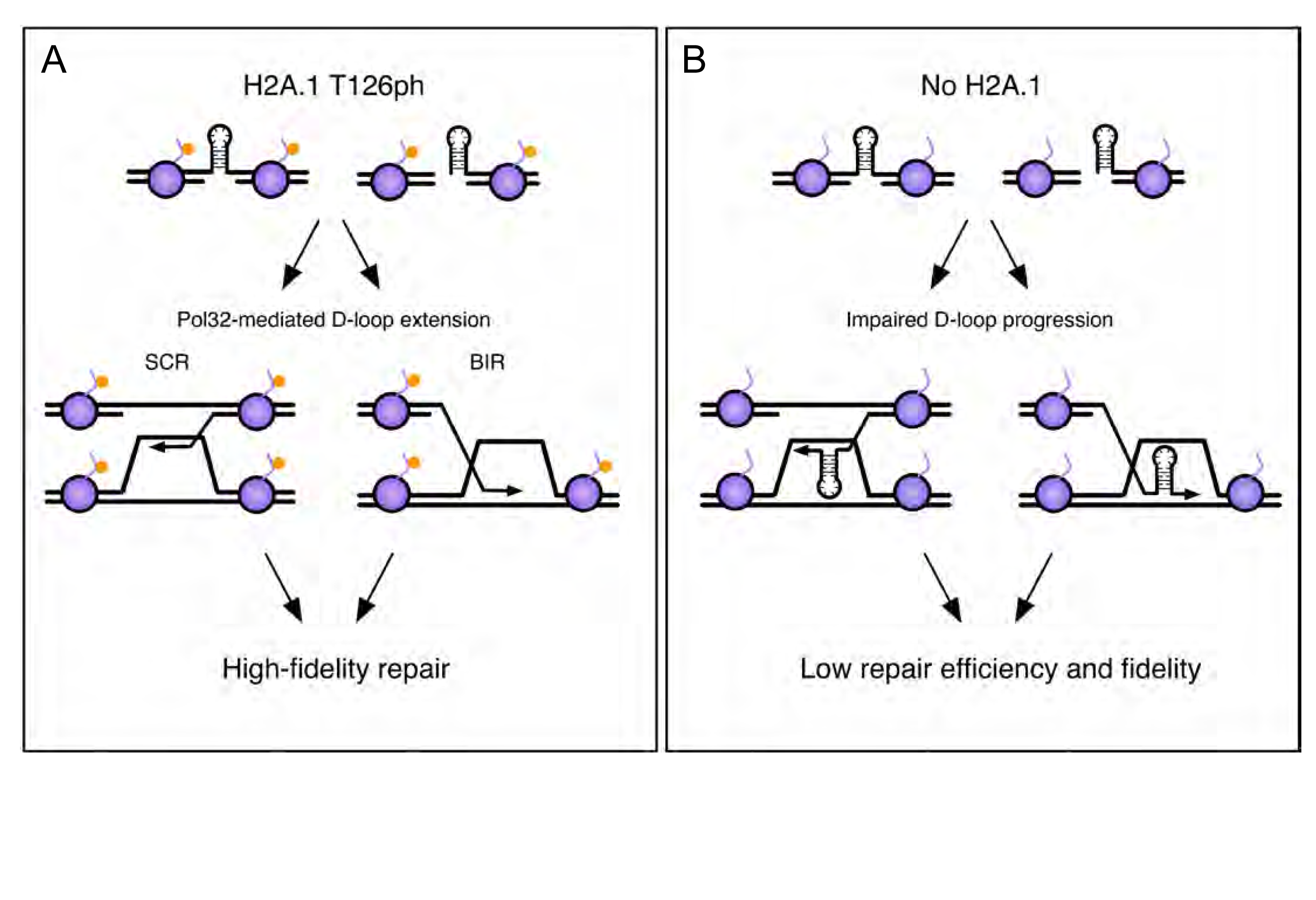
A model for high-fidelity HR-mediated repair promoted by H2A.1 and T126 phosphorylation. The initiating lesion at the CAG repeat could be a nick, gap, or a one-ended break (e.g. created at a replication fork). In the presence of a modifiable Thr126 residue (left pathway), phosphorylation will promote efficient D-loop extension after invasion into the sister chromatid, leading to repair with fidelity and maintenance of CAG repeat number. Alternatively, the resolution of recombination intermediates may require Thr126ph to prevent multiple reinvasion events that could lead to repeat expansions (not shown). Without efficient Thr126 phosphorylation (right pathway), or phosphorylation at the less efficient Thr125 residue in H2A.2, D-loop extension is impeded, allowing hairpins or misalignment during Rad51-dependent invasion, leading to less efficient repair and CAG repeat instability.

Perhaps less commonly, a replication fork stalled by a CAG or CTG hairpin may be converted to a one-ended DSB that could facilitate HR-dependent fork restart, similar to BIR (e.g. broken fork repair or BFR (46)). If fork restart proceeds with low fidelity, such as if recombination is misaligned or hairpins cause strand slippage, mutations can arise. Our data suggest that H2A.1 is required for efficient BIR, and CAG expansions occurring in the *hta1Δ* background were reduced when Pol32 was deleted, inhibiting BIR. Indeed, recovery from broken forks via BIR/BFR has recently been proposed to lead to large-scale expansions of a (CAG)_140_ repeat tract in yeast (55), and our data at the (CAG)_85_ repeat are consistent with these results. However, in the wild-type background, expansion frequency is significantly increased in a *pol32Δ* mutant where BIR is suppressed (Figure 5B), indicating that BIR is not the only pathway creating expansions. It does appear that, at least in part, that expansions in the *pol32*Δ mutant occur during recombination, suggesting that efficient D-loop synthesis through a CAG repeat requires Pol32 (Figure 5B). However, it is possible that some expansions are also arising by a different mechanism, such as impaired Polδ synthesis during replication.

## Conclusions

At the occurrence of DNA damage, recombination is thought to be more protective to genome integrity than end joining because the repair is templated. However, recombination can itself be mutagenic if it does not proceed in a regulated manner (22, 56). A turnover of chromatin modifications during repair is an attractive model for facilitating proper repair progression, either by influencing chromatin reorganization (reviewed in (57)) or by facilitating sequential recruitment of repair factors (reviewed in (1, 3)). Although the density of nucleosomes can vary at different genomic loci, the abundance of histones throughout the genome means they are readily available at the frontlines of DNA damage. In yeast, histone H2A.1 and H2A.2 are both present in a healthy cell and are likely interspersed throughout the genome. Transcript from H2A.2 is present in higher proportion than H2A.1 under normal conditions (unpublished data cited in (29)), but there must be an adequate amount of H2A.1 present on the chromatin to act in a fashion that promotes DNA repair. This may be similar to H2AX distribution in human cells, which is 2.5-25% of the total H2A pool (48). Further, in mammalian cells the H2AX histone variant is also required for efficient SCR and to prevent mutagenic single strand annealing of DSBs (58). The analogous H2AX threonine (T136) is also phosphorylated in mammalian cells (59, 60). Thus, it would be interesting to test whether mammalian T136 phosphorylation plays an analogous role in repair fidelity as the one described here for yeast H2A.1-T126.

Although we have not yet identified a direct interactor with H2A.1-T126ph, any repair factor that contains a BRCT, WW, of FHA domain is a potential candidate. While BRCT and WW protein domains can recognize both pThr and pSer residues, FHA domains appear to be more specific to pThr recognition (61). Therefore, the pattern of pThr126 and pSer129 modification could potentially modulate the binding of factors during repair. Depending on the phosphorylation status of both residues, a repair protein could recognize one or both residues, which could influence the overall progression of the repair process. Our results demonstrate a genetic interaction between H2A.1 and Pol32 (Polδ) in maintaining CAG repeat stability. Pol32 does not contain a phospho-binding domain and therefore a direct physical interaction between H2A.1 and Pol32 is unlikely, but they may interact via other protein mediators during repair. The timing and reading of H2A.1-T126ph and other chromatin marks may determine how effectively the Polδ complex moves through the donor strand during repair, ensuring that repair is efficient (timely) and that it proceeds with high-fidelity, limiting mutagenic repair outcomes. These will be interesting models to test in future experiments.

## Materials and Methods

### Yeast strain construction

The yeast strains used in this study are described in Table S7. *Mutant strain construction:* Genes of interest were deleted via PCR-based gene replacement with selectable gene markers. Transformants were grown on selective media; correct integration of the selectable marker was verified using PCR at the 3’ and 5’ integration junctions and primers internal to the target gene to verify ORF absence. *hta1 point mutants:* FY406 H2A point mutant strains were obtained from Dr. Jessica Downs (35). Both the TRT1 and TRT2 loci are deleted, thus the cell is lacking both the *HTA1/HTB1* and *HTA2/HTB2* loci. The H2A point mutants were introduced on a CEN plasmid containing either *hta1-T126A* or *hta1-S129A* point mutation, a wild-type copy of *HTB1*, and a HIS3 marker. A YAC containing a (CAG)_85_ repeat tract was introduced by mating with a karyogamy-deficient mutant strain (Kar1-1) that carries the YAC (CFY1736). For genomic integration of *hta1* point mutants, the *HTA1* gene containing either the T126A or S129A point mutants was PCR amplified from a CEN plasmid containing either *hta1-T126A* or *hta1-S129A* point mutation and cloned into the pFA6a-KanMX6 plasmid. The entire HTA1 gene and KANMX marker with a 40bp tail homologous to the 3’ end of HTA1 was PCR amplified and transformed into yeast. Integration of the point mutation was confirmed by PCR and sequencing. *Replacement of HTA2 in HTA1 gene locus*: A selectable KANMX6 gene was knocked-in 150 bp downstream of the HTA2 stop codon. The entire *HTA2* gene and KANMX6 marker was PCR amplified with primers with 40 bp tails homologous to the *HTA1* locus. Integration of *HTA2*+KANMX6 into the HTA1 gene locus was confirmed by PCR and sequencing.

### CAG repeat stability assays

PCR-based stability assays were performed as described (23, 24). Briefly, single colonies with verified (CAG)_85_ tract lengths (minimum four per strain) were inoculated in 2 ml YC-Leu or YC-Leu-Ura to OD_600_ of 0.03 to select for the YAC. Cells were grown for 6-8 generations to an OD_600_ of 1.9-2.8, at 30oC with rotation. A 10-4 dilution of cells was plated on YC-Leu-Ura to select for both arms of the YAC for H2A variant mutants, and YC-Leu-Ura-His for the H2A point mutant strains to select for the YAC and the presence of the plasmid. The CAG tract lengths of approximately 26 daughter colonies per mother colony inoculated were tested by whole colony PCR. PCR products were run in a 2% MetaPhor agarose gel or a custom gel mix on a fragment analyzer (Custom kit DNF945, Advanced Analytical Technologies, Inc), and repeat length was estimated within 2% accuracy by comparison to DNA ladders. The number of expansions and contractions were evaluated for significant deviation from wild-type cells using Fisher’s Exact Test (Table S2).

### Chromatin analysis by MNase digestion and Indirect End-labeling

Chromatin digestion was performed as previously described ((62), adapted from (63)), with the following modifications: spheroblasts were digested with 0, 0.25, 2.5, or 7.5 units of MNase and the DNA pellet RNase A digested for 30 minutes. The DNA was extracted twice using an equal volume of chloroform and finally precipitated by adding 67 μl 7.5M NH_4_OAc and 500 μl isopropanol. *Southern Detection:* MNase digested DNA (20-30 μg) was run on 1.5% agarose at 80V for 6hr and then Southern blotted as previously described (62). Chromatin structure was probed with a P^32^ labeled 358 bp PCR fragment amplified from 102 bp upstream of the CAG repeat on YAC CF1.

### Mononucleosome positioning detection by Illumina Bead Array

Chromatin was isolated and MNase digested as described above, except mononucleosomes were prepared by digesting the chromatin with 10 units of MNase for 15 minutes. Purified mononucleosomal DNA was amplified using the GenomePlex Whole Genome Amplification kit (Sigma). A sample was run in 1.5% agarose to verify amplification and to verify that fragments were ∼150 bp. Amplified mononuclesomes were purified with a GenElute PCR clean-up kit (Sigma) and subsequently 3’ biotin end-labled (Pierce); the DNA was chloroform extracted, and labeling was verified using a dot blot according to the manufacturer’s instructions. Purified mononucleosomes were applied to a custom Illumina array that contained YAC CF1 sequence spanning a region 425 bp upstream to 438 bp downstream of the repeat tract in 30 bp non-overlapping probes (Table S3). For each sample, a 12 μl aliquot of the purified mononucleosomal DNA was mixed with 3 μl of 15.3X SSC buffer containing 2.4% SDS, heated at 95^°^C for 5 minutes, and snap cooled. The sample was applied to the sample zone of the microarray, mixed by pipetting, and capillary action pulled the sample across the microarray face under a the protective plastic overlay. Hybridization in a humidified chamber was carried out overnight at 62^°^C. After hybridization, the clear plastic overlay was removed and the arrays were submerged in stringency wash (0.2% SDS and 2X SSC buffer), gently inverted, and then rocked gently on a rocking platform for 5 min. The slide was then submerged in 0.05X SSC buffer, gently inverted and then vigorously rocked for 1 min. To reduce non-specific binding of the stain, arrays were incubated in 40 mL of blocker casein in 1X PBS (Thermo) for 5 min. Arrays were then stained in the dark for 10 min in 5 ml blocker casein with 1 µg/ml streptavidin-Cy3 in PBS (Invitrogen). Chips were agitated in 40 mL of 0.05X SSC buffer on the rocking mixer for 1 min on maximum speed. The slide was then gently dried with compressed air and scanned with a BeadArray Reader (Illumina) using Direct Hybridization settings with a factor of 5. Signal intensity was exported to Excel via BeadScan.

### H2A sequence alignments

All DNA sequences were acquired from the Saccharomyces Genome Database (https://www.yeastgenome.org/) and aligned using SerialCloner or SnapGene. All protein sequences were acquired from Uniprot (http://www.uniprot.org/) and aligned by Clustal Omega (http://www.ebi.ac.uk/Tools/msa/clustalo/)

### Western blotting

Strains were grown in YPD at 30°C with agitation to log phase (OD_600_ = 1); cells were either untreated or exposed to 0.01% MMS for 2hrs at 30^°^C with agitation. Lysates were extracted according to (64) and https://research.fhcrc.org/gottschling/en/protocols/yeast-protocols/protein-prep.html, and Western blotted onto PVDF. Blots were probed with anti-H2A (Active Motif; 39945; 1:5000), anti-H2A-T126ph (Aves Lab custom antibody; gift from Krebs lab; 1:2500), or anti-H2A-S129ph (Abcam; ab15083; 1:2500) in 2.5% milk in 1X PBS (pH 7.4). The signals were detected with HRP-conjugated secondary antibody (1:2500) and ECL (Pierce). Western blot signals were quantified by ImageJ. The fold change in signal from wt was determined by comparing the relative quantification value (rqv) of a mutant to the relative quantification value of the wt (rqv= ratio of the indicated antibody to the loading control, with background subtracted).

### Antibody information

The anti-H2A-T126ph custom chicken antibody was generated by Aves Lab (http://aveslab.com). The antibody was raised against the phosphopeptide KKSAKA[pT]KASQEL and validated by peptide dot blot (Fig. S3). The anti-H2A-T126ph antibody specifically recognized the pT126 peptide and not unphosphorylated T126 peptide. Additionally, it did not recognize either unmodified or phosphorylated S122 or S129 (Figure S3)(37). A second antibody created by the same procedure was not specific, thus the specificity of antibodies created in this manner is variable.

### Sister chromatid recombination assays

Assays were performed as previously described (23, 38). Briefly, Trp+ Ade-cells were resuspended in 5 ml YEPD and grown to saturation at 30^°^C. Total viable cell count was measured by plating 10^−5^ dilutions on yeast complete (YC) media and recombinants were selected by plating 10^−2^ dilutions on YC-Trp-Ade. Recombination rates were calculated by the method of the median and rates were tested for statistical deviation using the Student’s t-test (Table S2).

### Ectopic gene conversion assays

Assays were performed as previously described (43). Briefly, colonies on YEPD were resuspended in 3 ml YP+Lactate and grown for 2-3 divisions at 30^°^C with rotation. Cultures were serially diluted to 10^−4^ and plated in duplicate on YEPD and YEP+Galactose. Gene conversion rates were calculated as percent viability (number of colonies on YEP+Galactose divided by the number of colonies on YEPD).

### Break-induced recombination assays

Assays were modified from (47). Briefly, colonies on YEPD+Nourseothricin were picked, serially diluted and plated on YEPD and YEP+Galactose in duplicate. Colonies were counted and percent viability determined (number of colonies on YEP+Galactose divided by the number of colonies on YEPD). To determine the frequency of BIR and other types of repair, all YEP+Galactose colonies were pinned onto YEPD, YEPD+Nourseothricin, and YC-URA. BIR rate was calculated as number of URA+ NAT-cells divided by the number of colonies on YEPD.

## Acknowledgments

Thank you to J. Downs for the H2A point mutant strains, D.R. Walt for the customized Illumina BeadArray, L. Symington for the SCR assay system, R. Anand and J. Haber for the BIR and GC assay systems, Jason Shaffer for help remaking the *hta1Δ* mutant, and Danae Schulz for performing the intial screen that identified H2A.1 as a candidate for further study.

---

**Figure S1.**
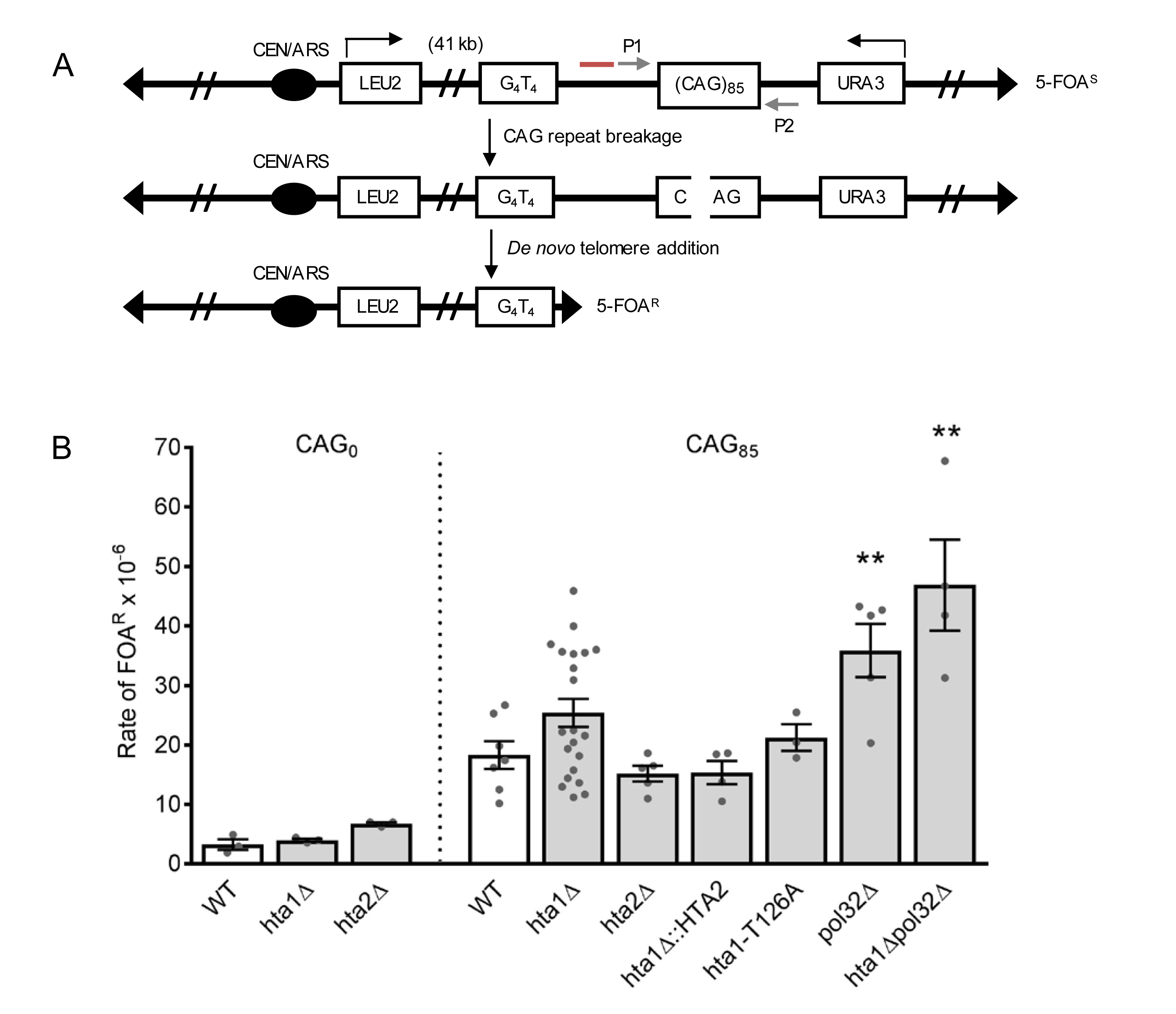
YAC system to evaluate CAG repeat fragility and instability. A) YAC CF1 contains an expanded CAG/CTG repeat originally cloned from a myotonic dystrophy patient. Here we have used 85 repeats with the CAG strand on the lagging strand template – referred to as (CAG)_85_. Breakage at the CAG repeat can result in resection to the backup telomere seed sequence (G_4_T_4_)_13_, resulting in *de novo* telomere addition and loss of the URA3 gene. This renders the cells resistant to 5-FOA and this rate can be used as a measure of repeat fragility (24). Primers (P1 and P2; gray arrows) flank the repeat to screen changes in repeat length by a PCR-based instability assay. Red line denotes location of Southern probe (described in Figure 2A). B) Fragility of YAC CF1 (either contained (CAG)_0_ (no tract) or a (CAG)_85_ repeat tract) was assessed as previously described (Reviewed in (24)). Briefly, single colonies with verified (CAG)_85_ tract lengths were incoculated were grown 6-8 divisions in YC-Leu liquid media at 30^°^C. To select for loss of the URA3 marker, a 10^−1^ dilution was plated on FOA-Leu media. Mutation frequency was calculated by the Method of Maximum Liklihood (FALCOR) (65). Rates were evaluated for significant deviations (as indicated) by the Student’s t-test, *p<0.05. Values and standard errors are listed in Table S1.

**Figure S2.**
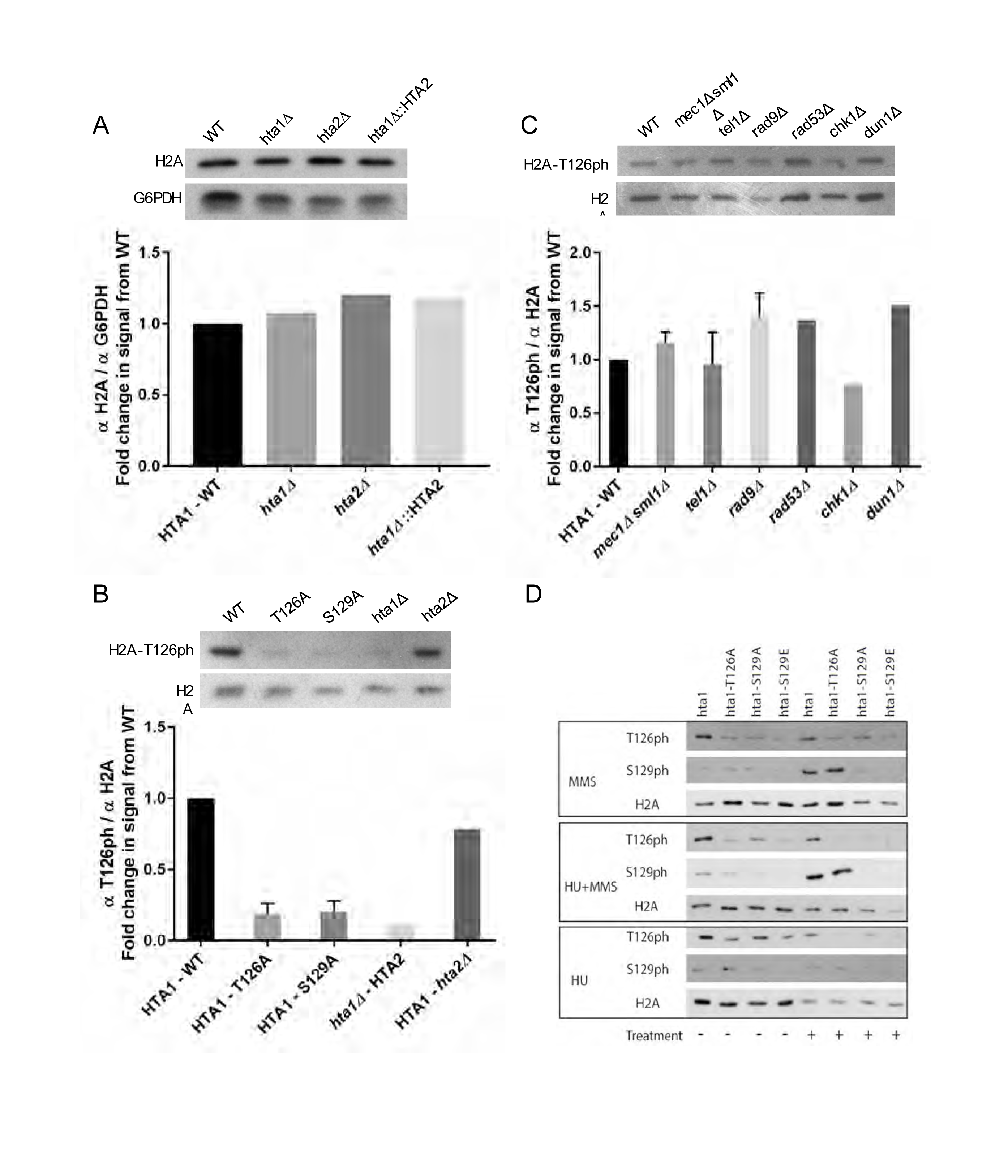
Western blot analyses of H2A and H2A phosphorylation levels. A) Total H2A levels in the H2A variant deletion mutants and replacement of H2A.1 with H2A.2. G6PDH was used as a loading control. The y-axis label, fold change in signal from wt, is the fold change in relative quantification values of the mutants from wt, where wt is 1. B) H2A-T126ph levels in the point mutant strains and full H2A copy deletion strains, using total H2A levels as a loading control. C) H2A-T126ph levels in kinase deletion mutants, using total H2A levels as a loading control. D) H2A-T126ph and H2A-S129ph levels in response to 0.03% MMS (top panel), 0.2M HU (bottom panel), or 0.03% MMS + 0.2M HU (middle panel).

**Figure S3.**
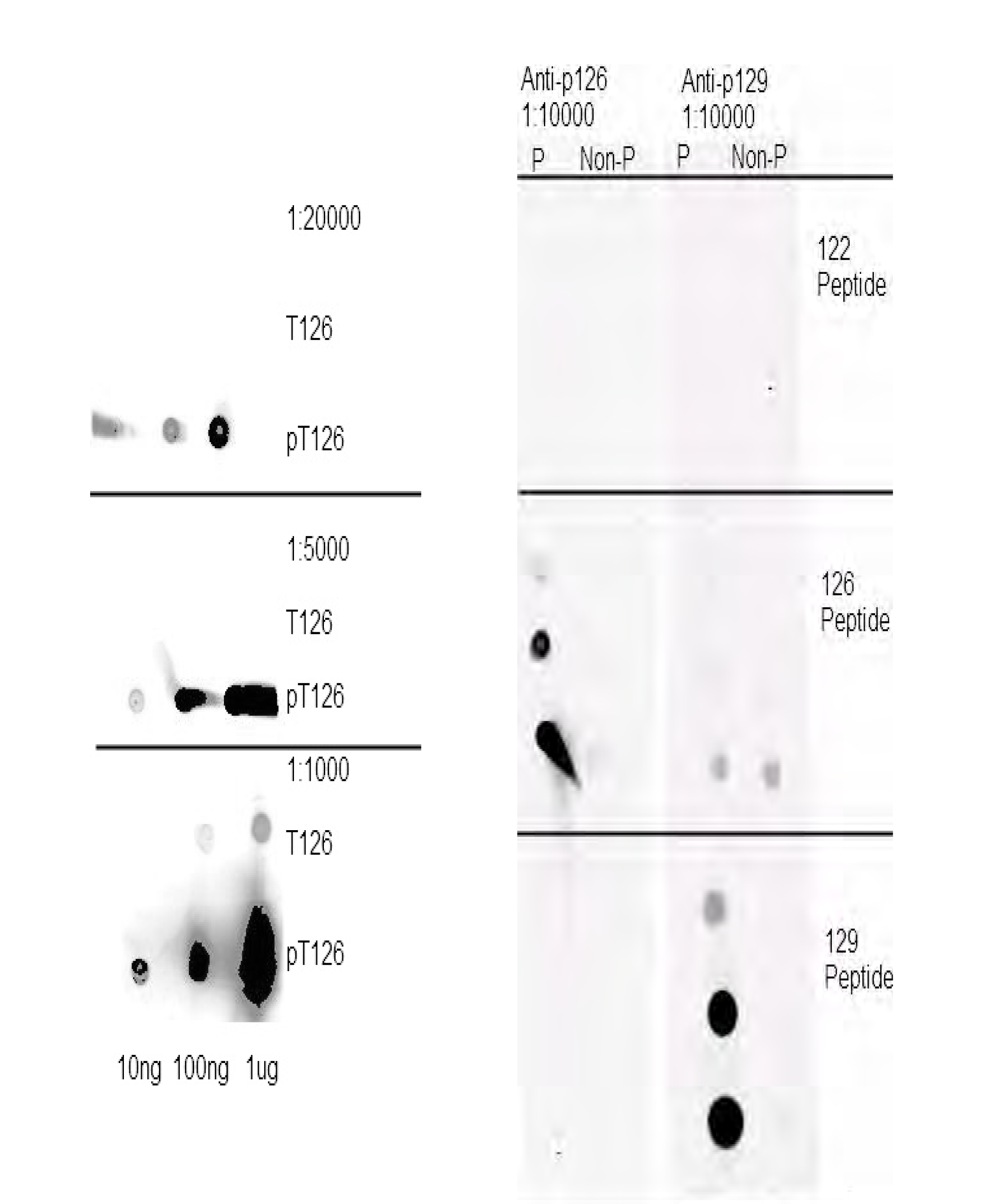
H2A.1 T126ph antibody dot blot. As originally appeared in (37). The H2A-T126ph antibody was raised against the phosphopeptide CZKKSAKA9pT]KASQEL. The antibody specificity was tested against H2A C-terminal tail phophopeptides, as indicated.

**Table S1.** (CAG)_0_ and (CAG)_85_ fragility assay data

|  | Strain |  |  |  |  |  |  |  |  |
| --- | --- | --- | --- | --- | --- | --- | --- | --- | --- |
|  | WT CAG-0 | *hta1Δ* CAG<sub>0</sub> | *hta2Δ* CAG<sub>0</sub> | WT CAG<sub>85</sub> | *hta1Δ* CAG<sub>85</sub> | *hta2Δ* CAG<sub>85</sub> | *hta1Δ::HTA2* *hta2Δ* CAG<sub>85</sub> | *hta1Δpol32Δ* CAG<sub>85</sub> | *pol32Δ* CAG<sub>85</sub> |
| Mean Rate<br>(x10<sup>-6</sup>) | **3.2** | **4.0** | **5.2** | **18.3** | **25.4** | **15.2** | **15.3** | **46.9\*\*** | **35.5\*\*** |
| SEM | 0.88 | 0.25 | 1.55 | 2.32 | 2.35 | 1.31 | 1.96 | 7.67 | 4.47 |
| No. replicates | (3) | (3) | (4) | (7) | (21) | (5) | (4) | (4) | (5) |

\*p<0.05 to wt, \*\*p<0.01 to wt, Student's t-test

**Table S2.**
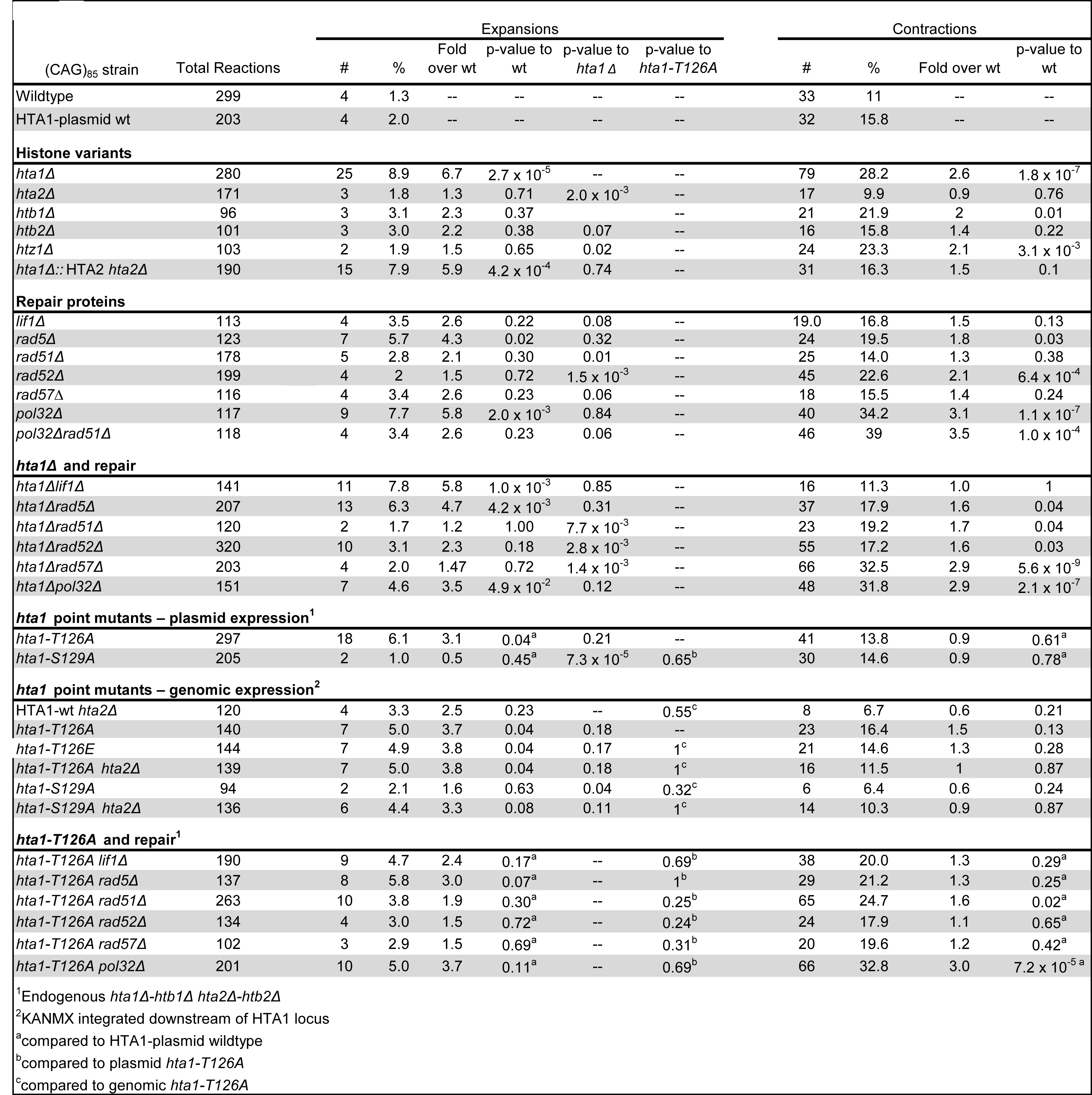
(CAG)_85_ repeat instability assay data.

**Table S3.**
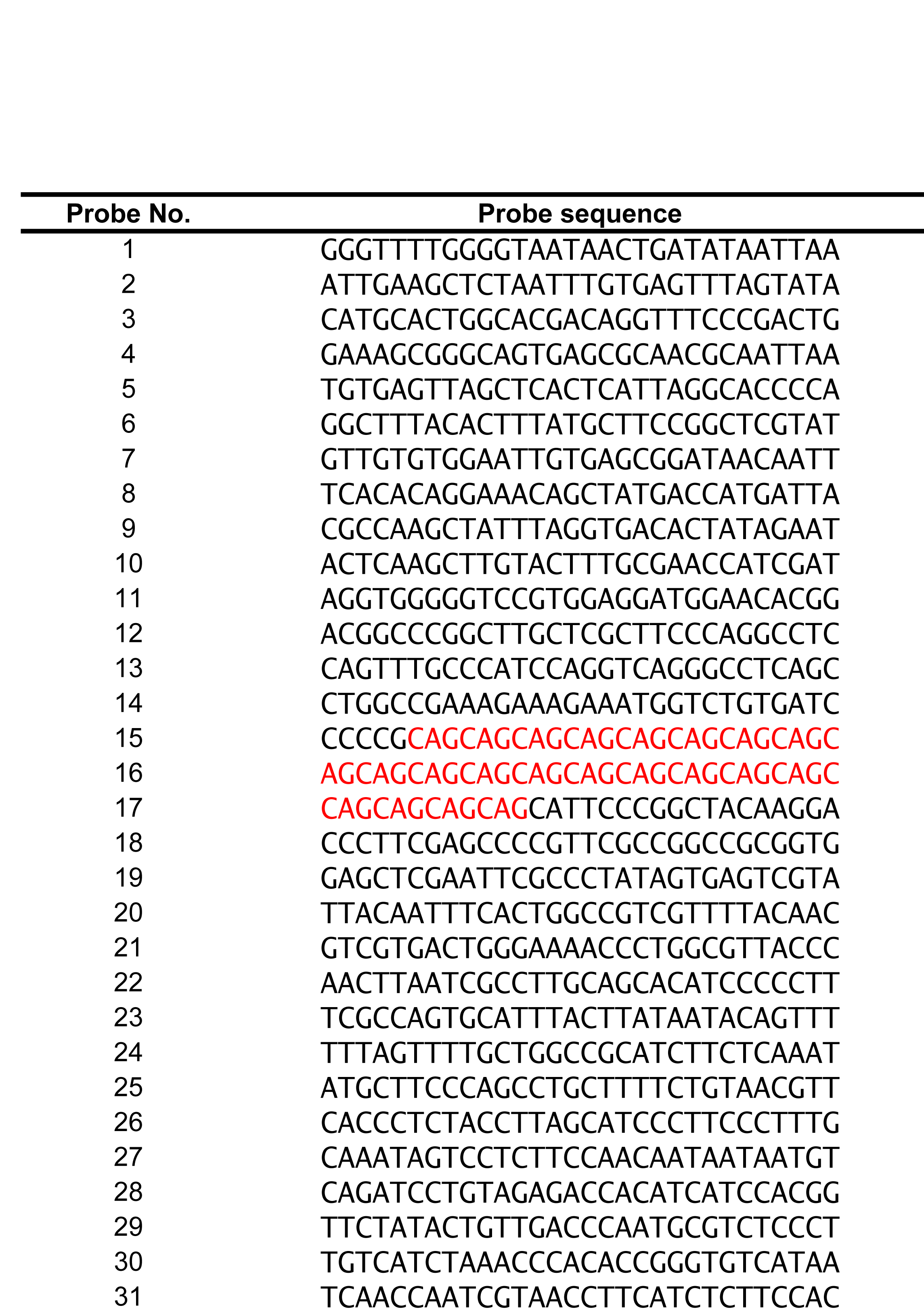
Illumina Array probe sequences. The 30-mer, non-overlapping probes spanned from 425 bp upstream of the repeat to 438 bp downstream of the repeat. The probes containing CAG repeats are highlighted. Probe 15 contains 8 repeat units, probe 16 contains 9 repeat units (pure repeats), probe 17 contains 4 repeat units. In total, the array contained 21 CAG repeats.

**Table S4.**
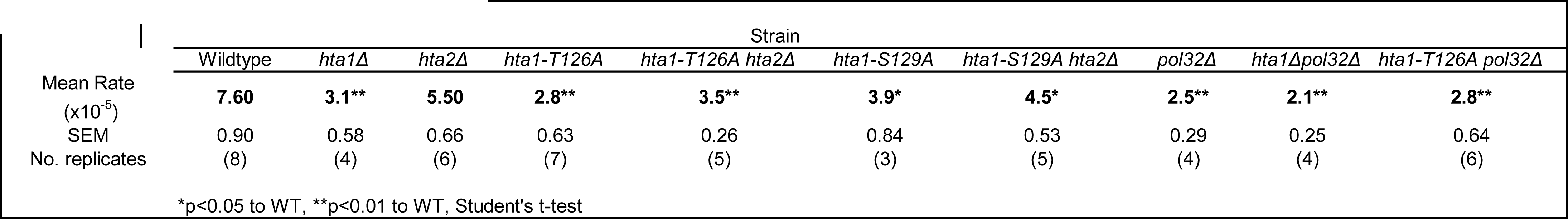
Sister chromatid recombination assay data

|  | Strain |  |  |  |  |  |  |  |  |  |  |  |
| --- | --- | --- | --- | --- | --- | --- | --- | --- | --- | --- | --- | --- |
|  | Wildtype | *hta1Δ* | *hta2Δ* | *hta1-T126A* | *hta1-T126A hta2Δ* | *hta1-S129A* | *hta1-S129A hta2Δ* | *pol32Δ* | *hta1Δpol32Δ* | *hta1-T126A pol32Δ* |  |  |
| Mean Rate<br>($\times 10^{-5}$) | **7.60** | **3.1\*\*** | **5.50** | **2.8\*\*** | **3.5\*\*** | **3.9\*** | **4.5\*** | **2.5\*\*** | **2.1\*\*** | **2.8\*\*** | | |
| SEM | 0.90 | 0.58 | 0.66 | 0.63 | 0.26 | 0.84 | 0.53 | 0.29 | 0.25 | 0.64 |  |  |
| No. replicates | (8) | (4) | (6) | (7) | (5) | (3) | (5) | (4) | (4) | (6) |  |  |
| *p<0.05 to WT, **p<0.01 to WT, Student's t-test |  |  |  |  |  |  |  |  |  |  |  |  |

**Table S5.** Ectopic gene conversion assay data

|  | Strain |  |  |
| --- | --- | --- | --- |
|  | Wildtype | *hta1Δ* | *hta2Δ* |
| Mean Viability (%) | **75.2** | **76.4** | **72.2** |
| SEM | 3.40 | 4.5 | 3.3 |
| No. replicates | (7) | (5) | (12) |
| *p<0.05 to WT, Student's t-test |  |  |  |

**Table S6.** Break-induced replication assay data.

|  |  | Strain |  |  |  |
| --- | --- | --- | --- | --- | --- |
|  |  | Wildtype | *hta1Δ* | *hta2Δ* | *pol32Δ* |
| BIR Rates | Mean Rate ($\times 10^{-2}$) | **7.60** | **3.1\*\*** | **5.50** | **2.5\*\*** |
|  | SEM | 0.90 | 0.58 | 0.66 | 0.29 |
|  | No. replicates | (8) | (4) | (6) | (4) |
| Repair Type | BIR (%) | 97.2 | 89.9 | 97.3 | 31.9 |
|  | NHEJ (%) | 0.9 | 1.6 | 0.6 | 35.2 |
|  | GC (%) | 0.3 | 4.8 | 0.4 | 12.5 |
|  | End loss/large deletion (%) | 1.7 | 3.9 | 1.7 | 20.4 |

\*p<0.05 to WT, \*\*p<0.01 to WT, Student's t-test

**Table S7.**
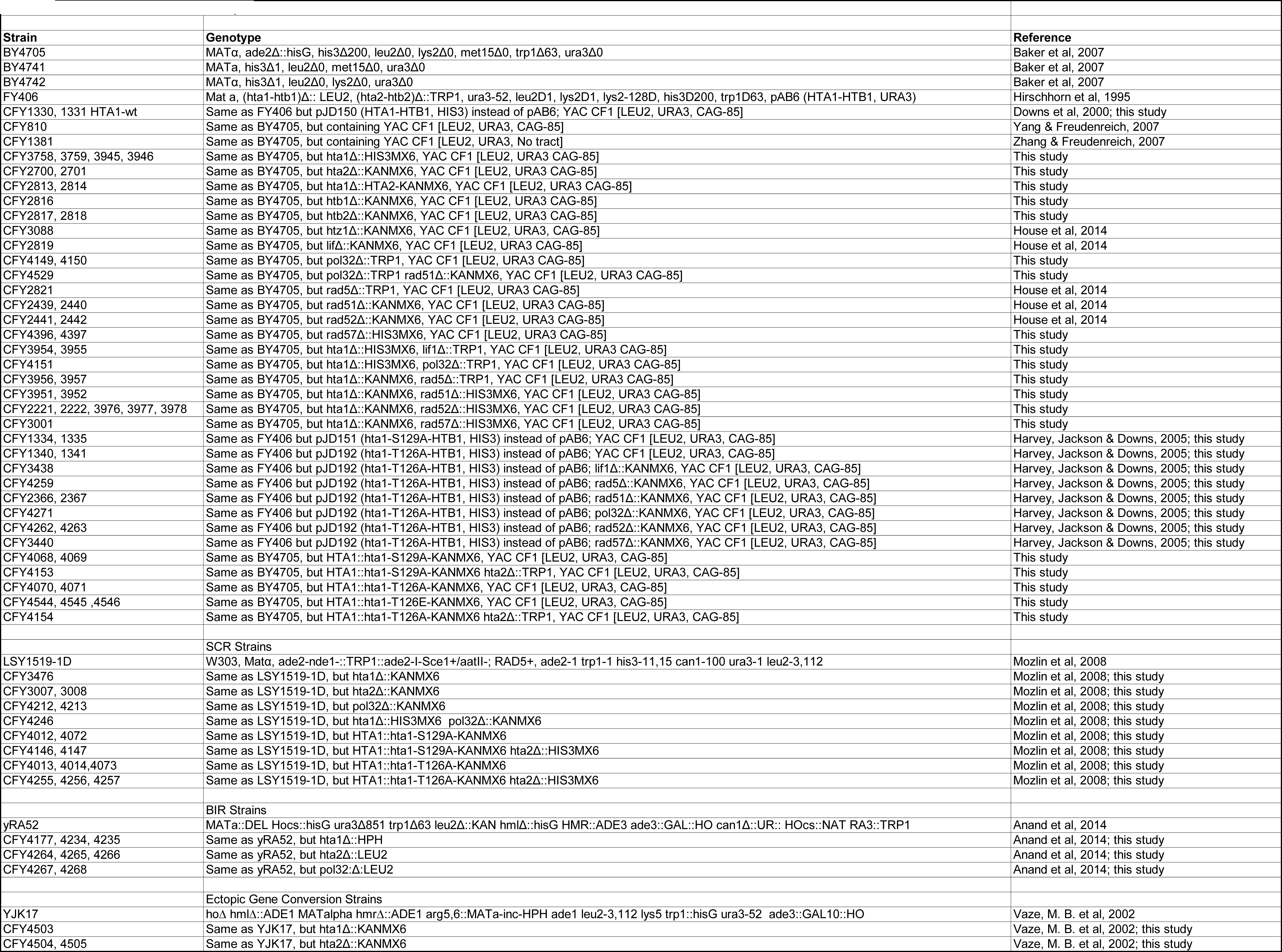
Yeast strains used in this study.

